# Fixed or random? On the reliability of mixed-effects models for a small number of levels in grouping variables

**DOI:** 10.1101/2021.05.03.442487

**Authors:** Johannes Oberpriller, Melina de Souza Leite, Maximilian Pichler

**Affiliations:** Theoretical Ecology, University of Regensburg, Universitätsstraße 31, 93053 Regensburg, Germany; Department of Ecology, University of São Paulo, Rua do Matão 321, Travessa 14, São Paulo, SP CEP 05508-090, Brazil

**Keywords:** Mixed-effects models, generalized linear models, multilevel models, hierarchical models, fixed effects, random effects

## Abstract

1. Biological data are often intrinsically hierarchical. Due to their ability to account for such dependencies, mixed-effects models have become a common analysis technique in ecology and evolution. While many questions around their theoretical foundations and practical applications are solved, one fundamental question is still highly debated: When facing a low number of levels should we model a grouping (blocking, clustering) variable as a random or fixed effect? In such situation, the variance of the random effect is imprecise, but whether this affects the statistical properties of the population effect is unclear.
2. Here, we analyzed the consequences of including a grouping variable as fixed or random effect in the correctly specified and other possible miss-specified models (too simple or too complex models) for data with small number of levels (2 - 8). For all these options, we calculated type I error rates and power. Moreover, we show how these statistical properties change with the study design.
3. We found that the model choice does not influence the statistical properties of the population effect when the effect is the same at all levels However, if an ecological effect differs among levels, using a random slope and intercept model, and switching to a fixed-effect model only in case of a singular fit, avoids overconfidence in the results. Additionally, power and type I error are strongly influenced by the number of and difference between levels.
4. We conclude that inferring the correct random effect structure is of high importance to get correct statistical properties. When in doubt, we recommend starting with the simpler model and using model diagnostics to identify missing components. When having identified the correct structure, we encourage to start with a mixed-effects model independent of the number of levels and switch to a fixed-effect model only in case of a singular fit. With these recommendations, we allow for more informative choices about study design and data analysis and thus make ecological inference with mixed-effects models more robust for small number of levels.

## Introduction

Many biological data from experimental or observational studies have a hierarchical grouping (or blocking or clustering) structure that introduces dependencies among observations (McMahon & Diez 2007; Bolker *et al*. 2009; Zuur *et al*. 2009; Harrison *et al*. 2018). When conducting statistical analyses, models have to reflect these dependencies in their assumptions for correct statistical properties (Arnqvist 2020), a task for which linear and generalized mixed-effects models (LMMs or GLMMs) were designed (Laird & Ware 1982; Chen & Dunson 2003). Mixed-effects models have also replaced ANOVAs as the common tool for variance analysis (Wainwright *et al*. 2007; Boisgontier & Cheval 2016) because they allow simultaneous analysis of variance at different hierarchical levels (Krueger & Tian 2004; Boisgontier & Cheval 2016), handle unbalanced study designs better (Swallow & Monahan 1984; Lindstrom & Bates 1988; Pinheiro & Bates 1995; Littell 2002), and have better statistical properties for missing data (Baayen *et al*. 2008). Therefore, mixed-effects models have become one of the most popular methods in ecology and evolution (Bolker *et al*. 2009) and other fields (e.g., psychology, see Meteyard & Davies 2020).

The ability of mixed-effects models to adapt to different data structures (i.e. their flexibility to handle many different hierarchical structures, see Box 1), which makes them in the first place so powerful (Wainwright *et al*. 2007), raises discussions about their proper application (Nakagawa & Schielzeth 2013; Dixon 2016). These issues include data-related problems such as overdispersion (Harrison 2014, 2015), robustness to wrong distributional assumptions (Schielzeth *et al*. 2020), and their correct evaluation (Nakagawa & Schielzeth 2013). Despite these rather technical challenges, there are also application-oriented issues (Harrison *et al*. 2017; Meteyard & Davies 2020) such as the question whether to use the most complex random effect structure (Barr *et al*. 2013; but see Matuschek *et al*. 2017), their correct interpretation (e.g. Dixon 2016), or in which situations we should model a grouping variable as random or fixed effect (Harrison *et al*. 2018).

A priori, modeling a grouping (or blocking) variable as fixed or random effect are equally well suited for multilevel analysis (Townsend *et al*. 2013; Kadane 2020) and strict rules don’t exist because the best strategy generally depends on the goal of the analysis (Gelman & Hill 2007, see Box 2). For instance, for unbalanced study designs, random effect estimates incorporate between and within group information whereas the corresponding fixed-effect model (grouping variable is specified as a fixed effect) only within group information(McLean *et al*. 1991; Dixon 2016; Shaver 2019; but see Giesselmann & Schmidt-Catran 2020). This is important when one is interested in the actual level effects themselves (also called narrow-sense inference analysis), but rather negligible when only interested in the population effect (broad-sense inference analysis), where the individual levels of the grouping variable are only nuisance parameters.

These inferential properties stem from different assumptions underlying these two options (specifying the grouping variable as fixed or random) (Millar & Anderson 2004). The most prominent difference is that modeling a grouping variable as random effect implicitly assumes that the individual levels of the grouping variable are realizations of a common distribution, usually a normal distribution, for which the variance and the mean (the population effect) are unknown and need to be estimated (e.g. DerSimonian & Laird 1986). This assumption shrinks the estimates of each random effect level to the mean of the underlying distribution. In contrast, modeling a grouping variable as a fixed effect makes no distributional assumptions about the individual level estimates (i.e., making them independent of each other and thus no between level information is used to estimate the effects). Thus, compared to when modeling the grouping variable as a fixed effect, the random effect model has to estimate a lower effective number of parameters (e.g. Gelman and Hill, 2005), which could lead to higher power to detect significant effects of population effects at the cost of a small bias towards the mean of the random effect estimates (e.g. Johnson et al. 2014).

Based on these properties, and if the grouping variable is only a nuisance parameter (broad-sense inference), random-effect modeling seems preferable over fixed-effect modeling. It is, however, unclear if these advantages remain when the number of levels is small (cf. also Harrison *et al*. 2018), because this might cause an imprecise and biased random effects’ variance estimate (Harrison, 2018), which then could influence the population effect estimate of the mixed-effects model (Hox *et al*. 2017).

The ecological literature suggests, as a rule of thumb, to only fit a grouping variable as random effect if it has a minimum of five, sometimes eight, levels to precisely estimate the random effect’ variance (Bolker 2015; Harrison 2015; Harrison *et al*. 2018). With four or fewer levels in the blocking variable, the preferred alternative is fitting it as a fixed effect (Gelman & Hill 2007; Bolker *et al*. 2009; Bolker 2015). In other disciplines other thresholds are proposed, e.g. 10-20 in psychology (McNeish & Stapleton, 2016) or 30-50 in sociology (Maas & Hox, 2005). To our knowledge, however, none of these values were based on a systematic analysis of how the choice of modeling the grouping variable affects the statistical estimates (i.e., type I error rate and power) of the population effects (i.e., the average slope or intercept over a grouping variable, henceforth called the ecological or population effect).

Here, we analyze a situation where a researcher wants to make inference about the population effect and draw conclusions for the population of levels that were sampled and thus has decided to use a mixed-effect model, but is confronted with a low number of levels. We simulated a hypothetical unbalanced study design on reproductive success and height of a plant on a temperature gradient (altitudinal gradient among different mountains, see Box 1) to compare empirical statistical properties with a varying number of levels, blocks, or experimental units (two to eight mountains, the nuisance parameter) and two different data-generating processes. To represent the challenge of an analyst to correctly specify the model structure and the consequences if not, we additionally tested miss-specified models (too complex or too simple versions of the fixed and mixed-effects models). To investigate the consequences of these modeling choices on the population effect, we compared: type I error rates (how often a non-existing temperature effect would be interpreted as existing) and statistical power (the rate at which a truly existing temperature effect would be significant). Based on our results and in the context of broad-sense inference, we give practical recommendations on when to include grouping variables as random effects and in which situations it might be more beneficial to fall back to specifying them as fixed effects.

### Box 1: Example of an ecological study design with grouping/blocking variables

#### Sampling design

We want to understand the population effect of temperature on the reproductive success of a plant that grows in mountains. We hypothesize that: (H1) increasing temperature (lower altitude) increases the probability of flowering (reproduction), and (H2) the height of flowering plants. To do so, we establish altitudinal transects in many mountains (nuisance variable) and collect information from a certain number of plants.

#### Problem

The transects are not in the same geographical alignment, the type of soil varies in each mountain, and the plants are genetically very distinct among populations. All these factors introduce differences among populations that are not exactly of our interest (given our hypotheses), but statistically, plants of the same mountain are non-independent observations. The mountains can be considered as grouping, blocking, control, or nuisance parameters.

#### Hypothesis 1

The reproductive success (flowering or not) of plants increases with temperature.

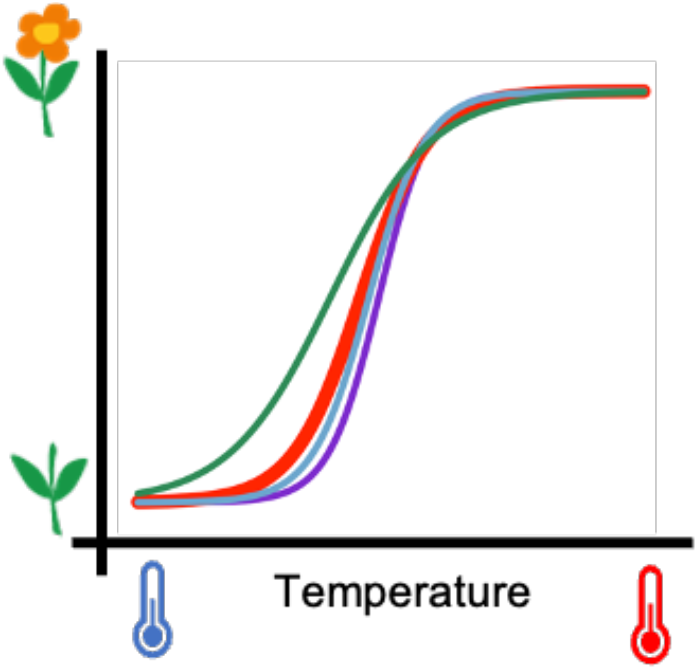

#### Sampling design

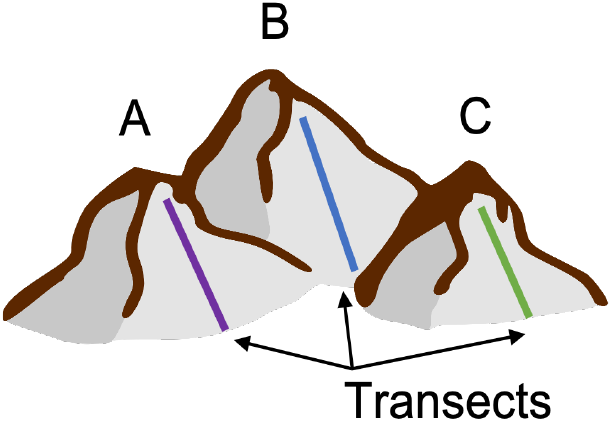

#### Modeling options

We may use a mixed-effects model with a random intercept and slope (Box 2) for mountain to account for the differences among populations (colored lines in H1 and H2), while still modeling the relationship of interest as fixed effects (red lines). An alternative may be to use a fixed-effects model, i.e., to include mountain as a categorical predictor (Box 2).

#### Hypothesis 2

The height of flowering plants increases with temperature.

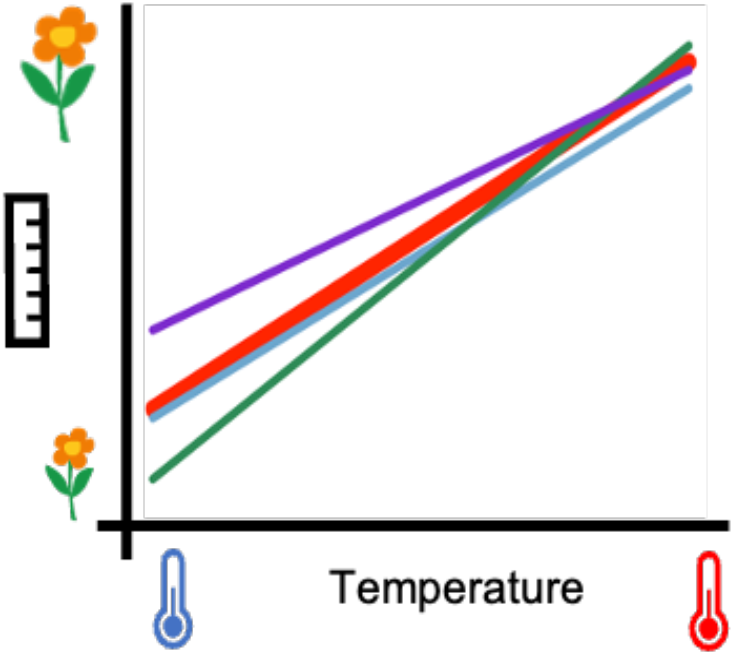

### Box 2: Differences between modeling a grouping variable as random or fixed effect

#### Fixed or random effects

The question of whether to include a hierarchical/blocking variable as random or fixed effect in the analysis depends on several factors. Fixed effects are usually used when the analysts are interested in the individual level estimates of a grouping variable (Bolker *et al*. 2009) and these are independent, mutually exclusive, and completely observed (e.g. control and treatment in experiments, male and female when analyzing differences between sex) (e.g. Hedges & Vevea 1998; Gunasekara *et al*. 2014). Random effects, in comparison, are modeling choices when the variance between the different levels (Bolker *et al*. 2009) and not the exact estimates of the different levels are of interest (e.g. DerSimonian & Laird 1986). Additionally, random effects can be used when not every realization of the underlying mechanism can be observed (e.g. species across a number of observational sites in different geographic areas) but the analysts want to control for its influence (i.e. pseudo-replication, see Arnqvist 2020). The two options differ in their interpretation, mixed-effect models use between- and within-group information whereas fixed-effect models use only within-group information. This subtle difference is important when for instance treatment or group differences are the goal of the analysis. Another important difference is that when modeling the categorical variable as fixed-effect conclusions apply to the levels used in the study, while when modeling as random-effect conclusions apply to the population of levels from where the studied levels were randomly sampled. However, in our example (Box 1), we are mainly interested in the population effect and not in the group differences which makes the inferential distinction negligible. See Gelman (2005) or Gelman & Hill 2007 for more decision criteria for whether an effect is random or fixed.

**Figure B1:**
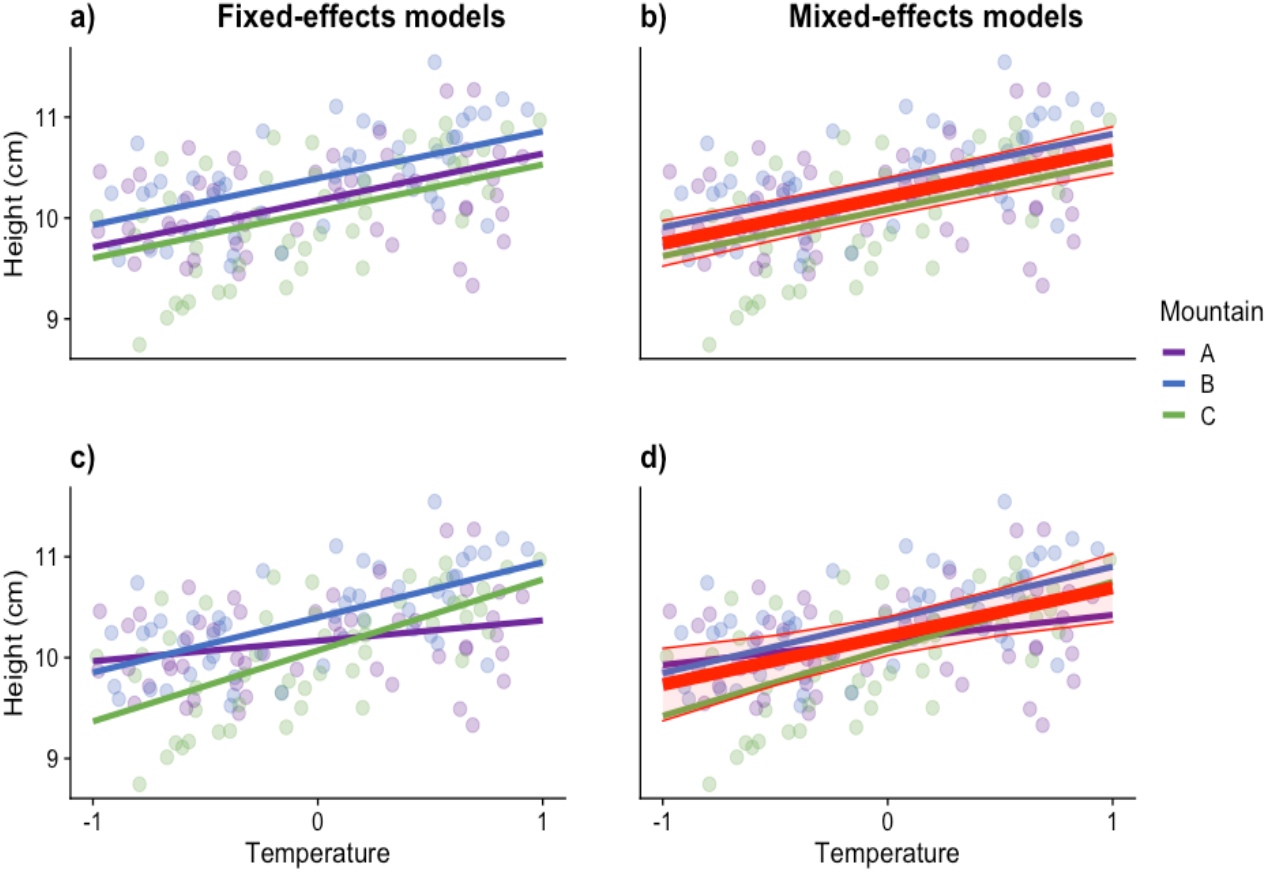
Fixed- and mixed-effects models fit to simulated data with random intercept (a,b) and random intercept and slope (c,d) for each mountain in the example from H2 Box 1. Dots represent the data points, lines the individual estimates for each mountain. The red line and shaded red intervals are the mean and its estimated standard deviation for mixed-effects models.

#### Technical differences between random and fixed effects

When specifying a grouping variable as fixed effect, the model with default contrast in R estimates the effect of one reference level (see Schielzeth 2010) differences between the reference level and possible linear combinations of other levels (Fig. B1a,c). Thus, fixed-effect models with default contrast don’t estimate mean effect over groups (i.e., the population effect), but it can be calculated using e.g. bootstrapping (see Supporting Information S1), with sum-to-zero contrasts, or follow-on packages such as emmeans (Lenth 2021). Mixed-effects models estimate the population effect and its standard deviation and from a Bayesian perspective each individual level effect or from a frequentist perspective predict future realizations of the individual random effect levels – Best Unbiased Linear Predictor (Fig. B1b, d). Blocking variables may not only imply different intercepts (Fig B1 a, b), but also different slopes (Fig B1 c, d - the temperature “ecological” effect). In fixed-effects models, this is done by introducing an interaction between the ecological effect and the grouping variable, which implicitly allows for correlations between slopes and intercepts. With mixed-effects models correlations between random slopes and random intercepts can be disabled. The choice of modeling different slopes and their correlation to intercepts for each group is related to the study design and may have impact on modeling structure and inference.

#### Models in R

Here are the formula R syntax1 of fixed and mixed-effects models fitted in Figure B1

*Scenario A* random intercept only

a) Height ∼ 0 + Temperature + Mountain (fixed-effect model ^2^)

b) Height ∼ Temperature + (1| Mountain) (mixed-effects model)

*Scenario B* random intercept and slope

c) Height ∼ 0 + Mountain + Temperature:Mountain (fixed-effect model^2^)

d) Height ∼ Temperature + (1|Mountain) + (0 + Temperature| Mountain) (mixed-effects model, uncorrelated random effects)

e) Height ∼ Temperature + (Temperature| Mountain). (mixed-effects model, correlated random effects)

1 From lm() or glm() base R functions, lme4 (Bates et al. 2015) and glmmTMB (Brooks et al. 2017) packages.

2 This formula parametrization estimates a separate intercept and slope (only in c) for each group independently. In c, the most common parametrization is *Mountain + Temperature + Mountain:Temperature*, which is the same as *Mountain*Temperature*, that uses one group as a reference intercept and slope and the other parameters estimate differences to the reference group. For purposes of prediction as in Fig B1 c, these parametrizations don’t change anything.

## Methods

### Simulated example

To compare random- and fixed-effect modeling of a grouping variable with small number of groups, we simulated data based on our hypothetical example from Box 1. We hypothesized first that higher temperatures improve the reproductive success (either yes or no) of a plant species (H1), and second, that higher temperatures also increase the average height of the reproductive plants (H2). To test these hypotheses, we simulated an unbalanced study design – a common scenario in ecology and evolution (Schielzeth et al. 2020) – with two to eight mountains from a varying number of plants for each mountain (H1: expected range between 40-360 plants per mountain, H2: expected range between 10-90 plants per mountain) while keeping the overall number of plants constant (H1: on average 200 plants per mountain, H2: on average 50 plants per mountain) along altitudinal transects.

We varied the number of mountains (groups) from two to eight and simulated 5,000 datasets for each case. Additionally, we performed a simulation, which varies components of the study design, to quantify their influence on the inference (see section Quantifying the influence of study design on type I error and power). We investigated performance for models with binomial (GLMM and GLM, H1) and normal distribution (LMM and LM, H2). The different choices of observations per group were made to meet the higher data request of binomial models compared to linear models.

### Scenarios of data complexity and model fitting

#### Scenario A - random intercepts per mountain

In scenario A, we assumed mountains only differ in their intercepts (mean height/reproductive success, Table 1, Eq. 1). When interested in the temperature effect, this is an example where the ecological effect is the same for each level of the grouping variable. For this scenario we tested two different mixed-effects model structures: a correctly specified model which corresponds to the data generating model (Table 1, Eq. 4) and a too complex model (Table 1, Eq. 5) with an additional random slope for each mountain. Since in real studies the true underlying data generating process is unknown, it is useful to understand if a too complex model can correctly estimate a zero (or nearly zero) random slope and, thus, approximate the true model structure (Table 1, Eq. 1).

**Table 1:**
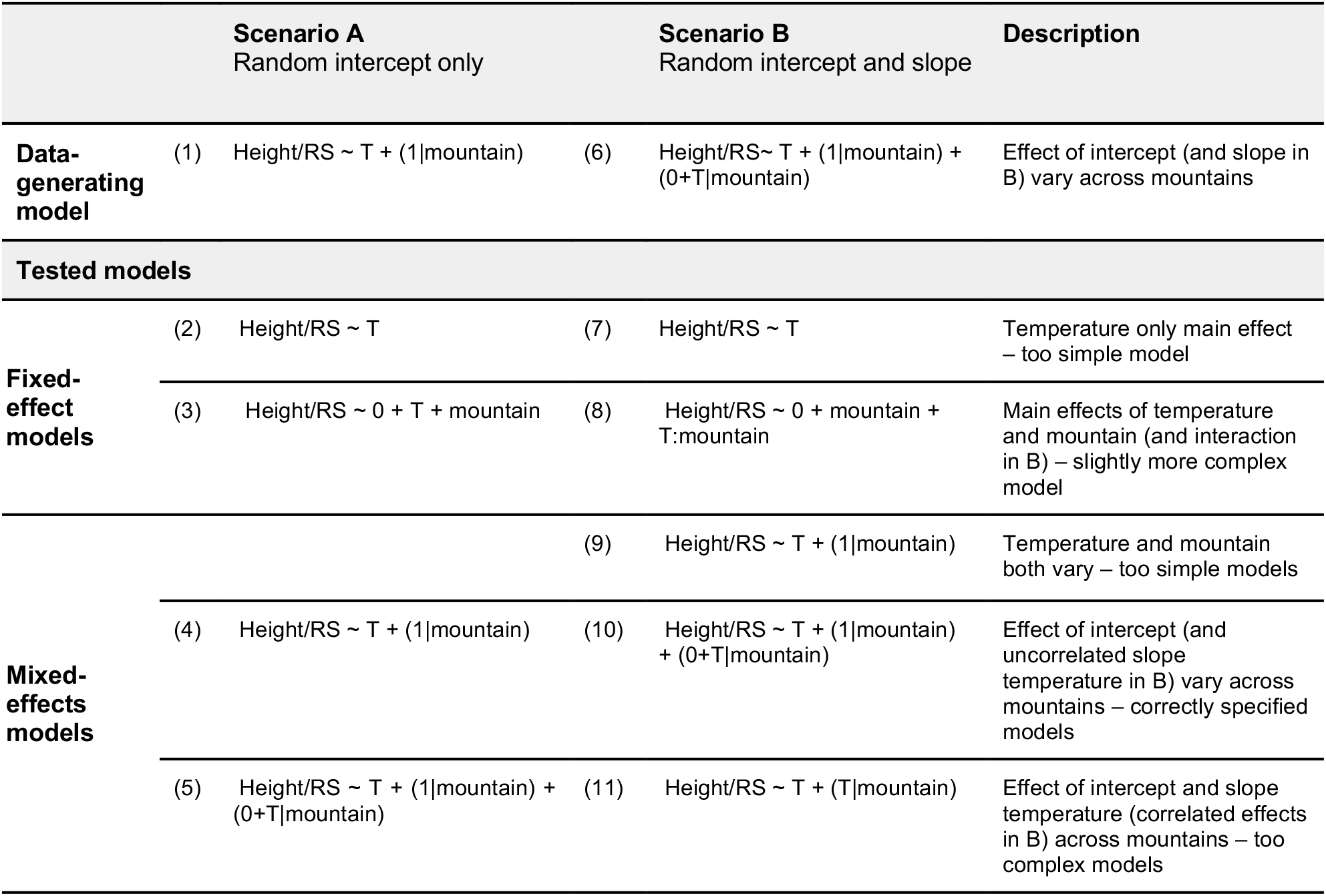
Data-generating and tested models for each scenario: Scenario A random intercept for each mountain and B random intercept and slope for each mountain. For the fixed-effects models, we used R syntax for model formula in *lm()* function and for the mixed-effects models we used syntax from *lmer()* or *glmmTMB()* functions from *lme4* and *glmmTMB* packages. Response variables are reproductive success (RS) for the binomial distribution models (H1, Box 1) and height of flowering plants for the normal distribution models (H2, Box 1). T is the temperature effect on both response variables.

As fixed-effect alternatives, we tested the correctly specified model with mountain as fixed intercept together with temperature as slope (Table 1, Eq. 3), and a too simple model omitting mountain at all (Table 1, Eq. 2). This last model corresponds to a mixed-effects model that estimates the random intercepts to zero.

#### Scenario B - random intercepts and random slopes per mountain

In scenario B, we assumed a random intercept and a random slope (without correlation among them) for each mountain as data-generating process (Table 1, Eq. 6). When interested in the temperature effect, this is an example where the ecological effect differs among levels of the grouping variable. We tested three different mixed-effects model structures: a correctly specified model corresponding to the data generating model (Table 1, Eq. 10), a too complex model with a correlation term for the random intercept and random slope (Table 1, Eq. 11), and a too simple model with only a random intercept for each mountain (Table 1, Eq. 9). We used the too simple model to test the effect of not accounting for important contributions to the data-generating process. Note, however, only in case of balanced designs and linear models the population effect estimate from the too simple model is consistent with the full model, because of different weighting schemes (for unbalanced designs) and the fact that the mean of a non-linear transformation (here link function) of estimates is not the same as the non-linear transformation of the mean of these estimates.

Specifying a correlation between the different random effects within a grouping-variable is the default in most mixed-effect model implementations, for a good reason, because omitting the correlation can lead to higher type I error rates (Matuschek *et al*. 2017). However, we disagree with the view of Matuschek *et al*. 2017 that trading a small increase in type I error rate for higher power is favorable, but it could still be an interesting solution with the often-small number of observations in ecological studies when the increase in power prevails the increase in type I error rate.

As fixed-effect alternatives, we tested the correctly specified model with the main effects of temperature and mountain and their interaction (Table 1, Eq. 8), and the too simple model dropping mountain as predictor (Table 1, Eq. 7). We tested the last model since mixed-effects models, with standard deviation estimates of zero for both random effects, correspond to fixed-effects models omitting the grouping variable.

### Model fitting

We fitted mixed-effects models to our simulated data with two of the most commonly used packages in R: *lme4* (Bates *et al*. 2015) and *glmmTMB* (Brooks *et al*. 2017). We present here *lme4* results because it reports singular fits, i.e., when some of the parameters of the variance-covariance Cholesky decomposition are exactly zero. Results with *glmmTMB* can be found in Supporting Information S1. To obtain p-values for mixed-effects models we used the *lmerTest* (Kuznetsova *et al*. 2017) package. For fixed-effects models in scenario A we extracted p-values from the summary and for scenario B we used the fitted variance-covariance matrix and the individual level effects to bootstrap the population effect and its standard error (see Supporting Information S1).

For LMMs we used the restricted maximum likelihood estimator (REML). For GLMMs we used the maximum likelihood estimator (MLE) as REML is not supported in *lme4* for GLMMs (for a comparison of REML and MLE see Supporting Information S1). All results of mixed-effect models presented in scenario A and B are for the datasets without singular fits (see section *Variances of random effects and singular fits*).

### Statistical properties and simulation setup

We compared the modeling options for both data generating scenarios based on important statistical properties of the temperature effect (ecological effect) in our simulated data: type I error rate and statistical power. Type I error rate is the probability to identify a temperature effect as statistically significant although the effect is zero. Statistical power is the probability to detect the temperature effect as significant if the effect is truly greater than zero. For a correctly calibrated statistical test, the type I error is expected to be equal to the alpha-level (in our case 5%), and the higher the statistical power the better the test. We additionally calculated coverage for part of the simulations, i.e., how often the true effect of temperature falls into the 95% confidence interval of the model, see Supporting Information S1 for results.

To investigate type I error rates for the slope/intercept, we simulated data with no slope/intercept effect, i.e., the effect of temperature/mountain on plant reproduction (average reproductive success of 0.4) and on height (average 0.4 cm) is zero. To additionally investigate statistical power and coverage, we simulated an example with a weak effect, i.e., an effect that is barely statistically significant. In our example, a weak effect corresponds to an average increase in size per unit step of the standardized temperature (linear scale) of 0.4 cm (Hypothesis 2) and 0.4 gain in reproductive success (Hypothesis 1) at scale of the linear predictors.

For scenario A and scenario B, the individual effects for each mountain were drawn from a normal distribution with standard deviation of 0.1 around the average effects: 0.4 cm average height (intercept), and 0.4 cm average increase in size or 0.4 (logit link scale) gain in reproductive success with temperature (slope).

### Variances of random effects and singular fits

To understand how the number of levels affected the random effect variance estimates, we recorded and compared them for random intercepts and slopes from the correctly specified mixed-effects model in scenario B (Table 1, Eq. 10). We also compared optimization routines (REML and MLE) in terms of estimating zero variances (singular fits, see below) for both LMMs and GLMMs (see Supporting Information S1). For bounded operations, which most R packages apply for the variance, one would expect a null distribution to be a combination of a point mass at zero and a chi-squared distribution (Stram & Lee 1994). For the sampling distribution with a true variance unequal to zero there are no proofs, but one would expect a similar distribution.

### Handling of singular fits

Technically, singular fits occur when at least one of the variances (diagonal elements) in the Cholesky decomposition of the variance-covariance matrix are exactly zero or correlations between different random effects are estimated as -1 or 1. However, how to deal with singular fits is being discussed. While Matuschek *et al*. 2017 suggests to think *a priori* about using simpler models, Barr *et al*. 2013 states to start with the maximal model and simplify the model in case of convergence issues and singular fits. While singular fits don’t signal a convergence issue, the common sense is that they should be prevented. When facing a singular fit, a practitioner usually changes the model structure and neglects the previous fit. Following this perspective, we decided to use only non-singular fits for calculating the above-mentioned statistical properties for the mixed-effect models. For fixed-effects models, we used estimates from non-singular and singular fits combined.

But using only non-singular fits for calculating the statistical properties impacts the statistical properties (e.g., type I error) because they are conditional on this selection and thus likely not to be at the nominal level (e.g., 5% for type I error rate). We calculated type I error rates for non-singular and singular fits combined which can be found in the Supporting Information S1 (Fig. S8, S10). For scenario B (random intercepts and random slopes) *lme4* gives only a generic singular fit warning but does not specify for which random effect the singular fit occurred (random intercept or random slope), which furthermore complicates the handling of singular fits (random intercept singular fit should be no problem for the population slope effect). But as our main intention is to report the type I error rates from the point of the practitioners who may adjust the model structure to dispose of the singular fit, our reported rates represent empirical type I error rates.

### Quantifying the influences of study design on power and type I error

The statistical properties of the ecological effect may depend not only on the number of levels (mountains) but also on the random effect’ variance, the overall number of observations and the balance of observations among levels. To further quantify their impacts on the statistical properties of the ecological effect, we additionally ran 1,000 iterations (each with 1,000 non-singular model fits) with the data generating model from scenario B for hypothesis 2 (height of plants, uncorrelated random slope and random intercept). Thereby, we sampled uniformly the number of mountains from 2 to 20, the random effects’ variance from 10^-4^ to 4, the overall number of observations from 10*number of mountains to 500*number of mountains. Additionally, to create different degrees of unbalancedness in data, we sampled for each mountain the average share of total observations from 0.1 to 0.9. Thereby, we ensured that there are at least 3 observations per mountain. We used the difference between the largest and the lowest proportion as proxy for the degree of unbalancedness.

We fitted the correctly specified LMMs and LMs from scenario B (Table 1, Eq 8) and calculated type I error rate and statistical power of the temperature effect. We then fitted a quantile regression using the *qgam* R-package (Fasiolo *et al*. 2020), with the statistical property (power and type I error rate) as response and variance, number of levels, total number of observations and the unbalancedness proxy as splines.

## Results

### Scenario A - random intercepts per mountain

When the ecological effect was the same among mountains, irrespectively of the number of levels (mountains), all models except for the too complex model (random intercept and slope) showed a type I error rate of 5% (Fig. 1a, c). Power increased (Fig. 1b) with the number of mountains for H2 from 90% (2 mountains) to 100% (5 to 8 mountains), and for H1 (Fig. 1d) from 35% (2 mountains) to 95% (8 mountains). Note that the model omitting the grouping variable presented similar properties as the other models. However, when increasing the standard deviation of the random intercept in the simulation, this model showed much lower power (Fig. S12).

**Figure 1:**
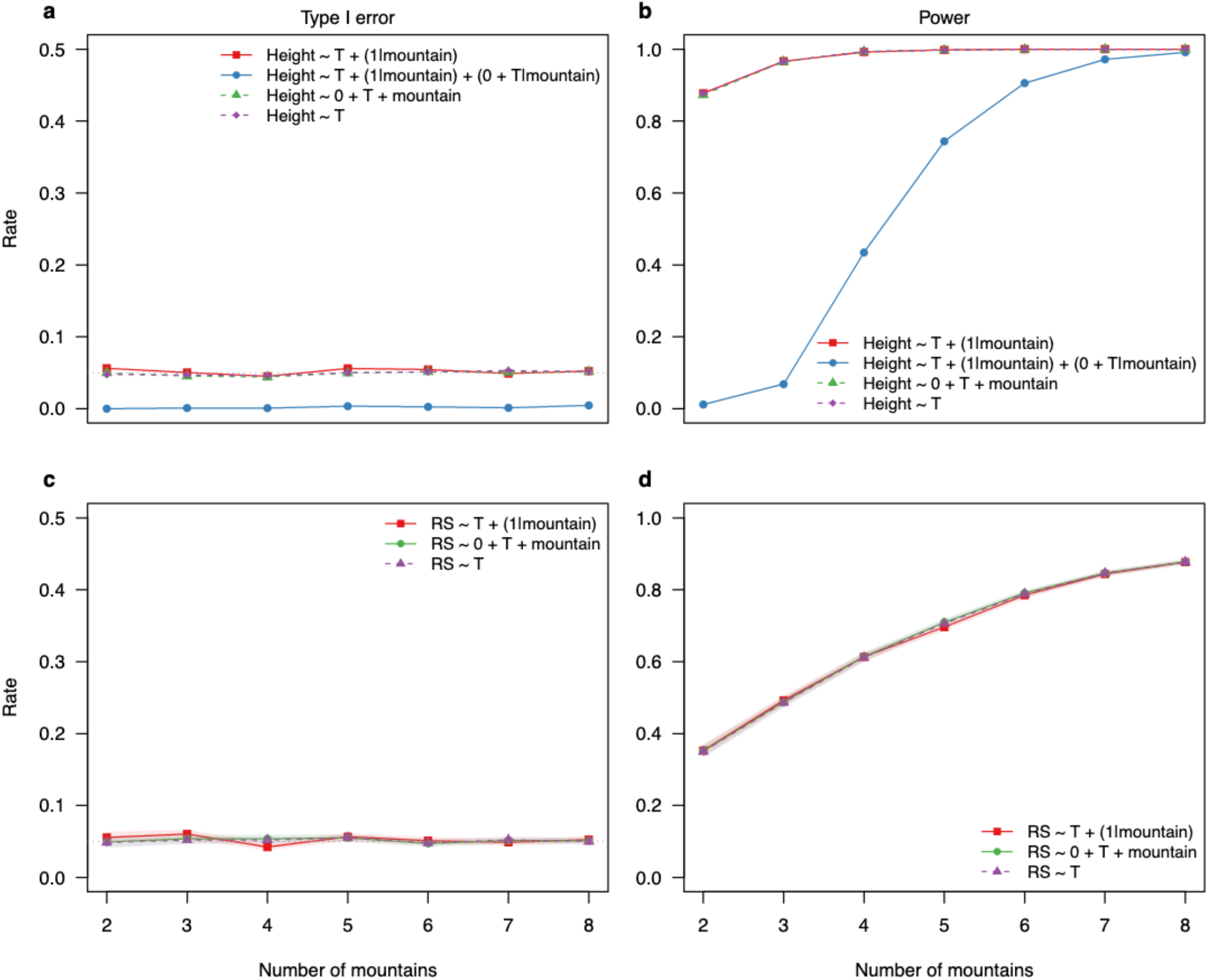
Type I error rates and power for linear fixed and mixed effect models (a, b) and generalized linear fixed and mixed-effects models (c, d), fitted to simulated data with 2-8 mountains (random intercept for each mountain - Scenario A) and 50 observations per mountain. For each scenario, 5,000 simulations and models were tested. Results for mixed-effects models are only from datasets in which mixed-effects models converged without presenting singular fit problems, while results for fixed-effects model are from all datasets.

For the too complex model for LMMs, we found on average a lower type I error rate of around 1-2% (Fig. 1a), and lower statistical power to detect the temperature effect for a small number of mountains (Fig. 1b). Binomial datasets with small number of observations per mountain (25, 50, 100) presented similar results regarding type I error but as expected, very low power among all models (Figure S14). The results for the intercept for the different models (see Fig. S9) are similar to the results for the slope in scenario B (see below).

### Scenario B - random intercepts and slopes per mountain

In scenario B, where the ecological effect differed among levels, the modeling decision had a more severe influence on the statistical properties (Fig. 2). We found that type I error rate of the correctly specified mixed-effects model (Table 1, Eq. 10) slightly increased (Fig. 2a, c) with the number of levels towards the nominal value (0.05) (Fig. 2a, c). The too complex model with correlated random intercept and random slope (Table 1, Eq. 11) presented similar properties, but with slightly increased type I error and decreased power (Fig. 2). For the correctly specified fixed-effect model, type I error (**≈** 2%) stayed constant with the number of levels (Fig. 2a, c) and power increased with the number of mountains. For H1 (normal model), however, the mixed-effects model showed higher power than the fixed-effects model irrespective of the number of mountains (Fig. 2 d). For both normal and binomial models, the too simple model omitting the grouping variable resulted in a higher type I error rate and higher power than the other models (Fig. 2). The results for the intercept are similar as the results for the slope (see Fig. S11).

**Figure 2:**
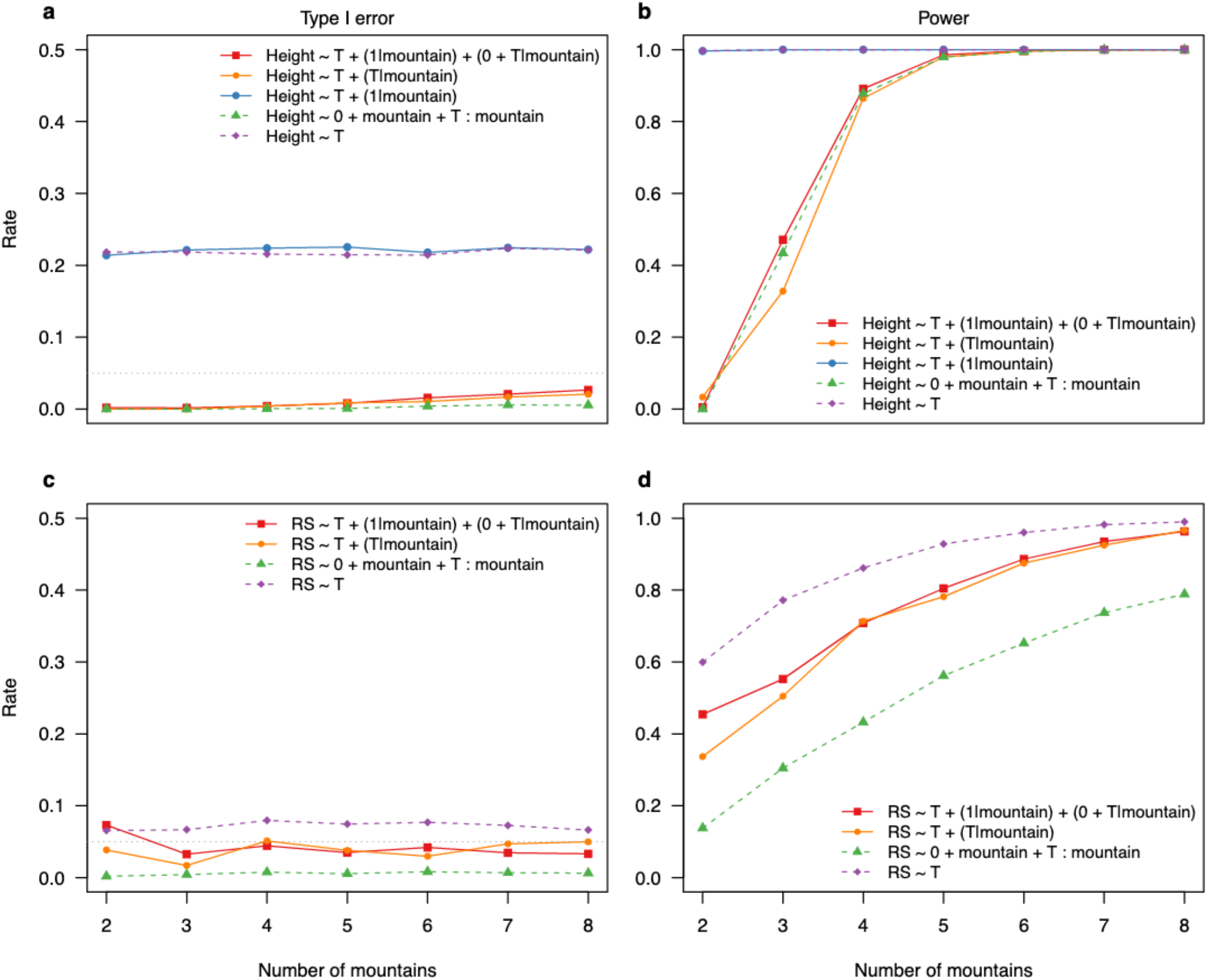
Type I error rates and power for linear (mixed effect) models (a, b) and generalized linear (mixed-effects) models (c, d), fitted to simulated data with 2-8 mountains for scenario B (random intercept and random slope for each mountain range). For each scenario, 5,000 simulations and models were tested. Results for mixed-effects models are only from datasets in which mixed-effects models converged without presenting singular fit problems, while results for fixed-effects model are from all datasets.

### Variance estimates of random effects and singular fits

We found for LMMs (singular and non-singular fit results combined) in Scenario B (random intercept and slope) that random effects’ variance estimates of the correctly specified model (Table 1, Eq. 10) showed bimodal distributions with a chi-squared distribution around the correct value (0.01) and a point mass at zero (Fig. 3a, b median is near to zero). The point mass at zero decreased in height with increasing levels, i.e., less models estimated a variance of zero with an increasing number of mountains (Fig. 3a, b, see also Table S1). There was smaller bias for the random intercept variance estimates than for the random slope variance estimates, which were still biased for eight levels. When looking at models without singular fits, the variance estimates were chi-square distributed (Fig. 3c, d). The bias towards higher values was stronger compared to estimates with singular fits, especially for the random slope estimates (Fig. 3d). Results for *glmmTMB* were consistent with *lme4* when using a threshold of 10^-3^ to classify a variance as zero (singular fit) (Fig. S2).

**Figure 3:**
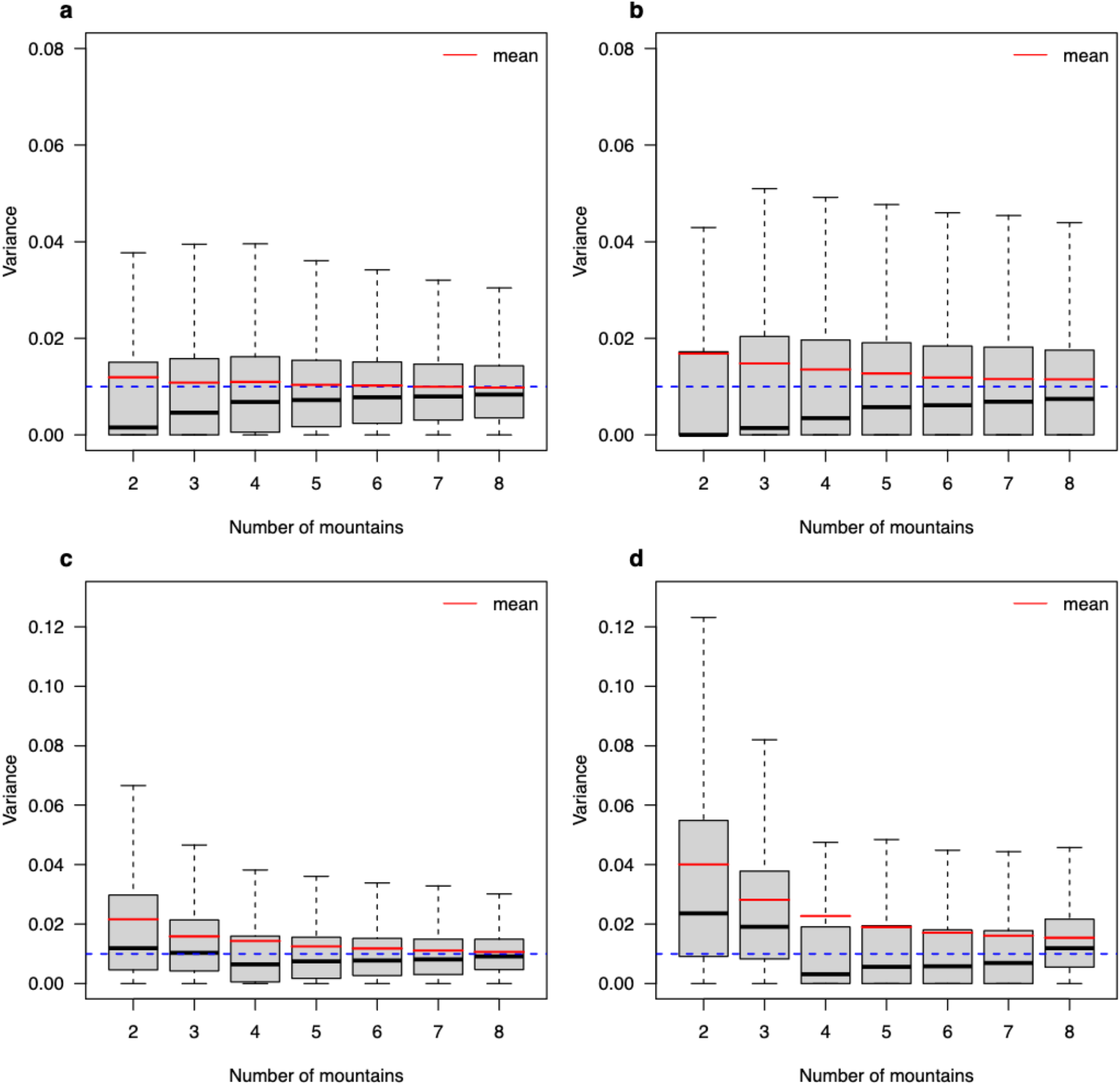
Variance estimates of random intercepts (a, c) and random slopes (b, d) for linear mixed-effects models (LMM, Table 1. eq. 10) in Scenario B, fitted with *lme4* using REML to simulated data with 2-8 mountains. Figures (a) and (b) show the results for all models (singular and non-singular fits) and figures (c) and (d) show the results for only non-singular fits. For each scenario, 5.000 simulations and models were tested. The blue dotted lines represent the true variance used in the simulation (0.01) and the red lines the average variance estimates.

By comparing the fitting algorithms, we found that using MLE led to more zero-variance estimates, i.e., singular fits, (Fig. S3, S4) than REML. Additionally, using MLE, also non-singular variance estimates were strongly biased (Fig. S3, S4), but the bias decreases with increasing number of levels. For both optimization routines, increasing the number of levels reduced the number of singular fits (Table S1).

We found that singular fits had a high influence on type I error and power (Fig. 4) of mixed- and fixed-effects models. For singular fits, the type I error rate of the correctly specified mixed-effects model was constant around 10% (similar to the model omitting the grouping variable), while with non-singular fits it was 1% for two levels and increased towards 3% with eight levels (Fig. 4a). In comparison, the fixed-effects model had similar type I error rates singular and non-singular fits, both increasing from 0% (two levels) towards 1% (eight levels) (Fig. 4c).

**Figure 4:**
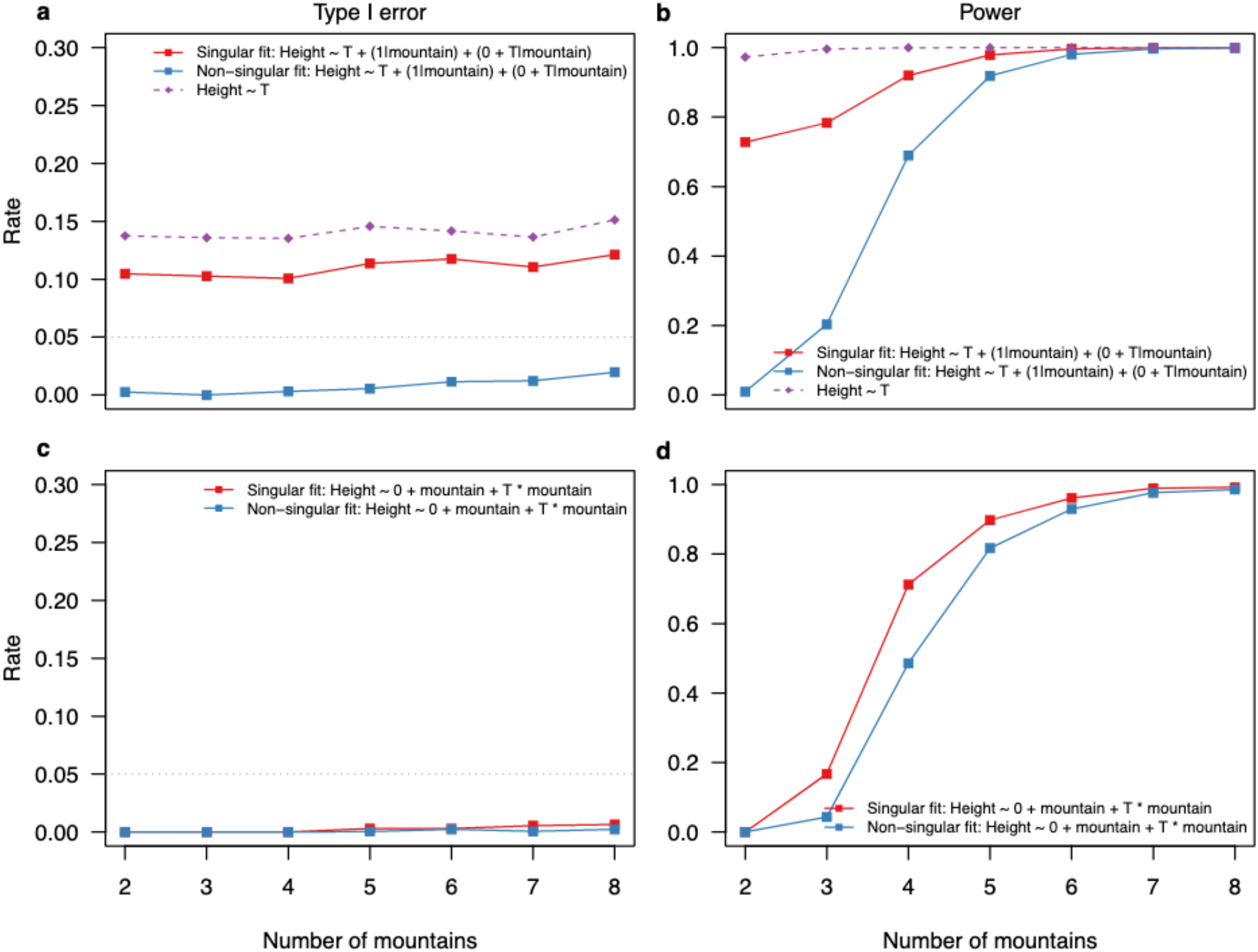
Type I error rate and power of the correctly specified linear fixed and mixed-effects models in scenario B. We separated the datasets based on if when fitted with lme4 they presented a singular fit (red lines) or non-singular fit (blue lines). Figure (a) and (b) are results for the linear mixed-effects models, and (c) and (d) for the linear fixed-effects models. For comparisons, we show also results for the fixed-effects model that omits the grouping variable (mountain).

We also found differences in power for the models between singular and non-singular fits (Fig. 4b, d). The power of the mixed-effects model with correct structure is higher for singular than non-singular fits especially for a low number of mountains (Fig. 4b). For the fixed-effects model, this difference is less strong, however the model has still higher power for singular than for non-singular fits (Fig. 4d).

### Quantifying the influences of study design on power and type I error

We found that the average type I error of mixed-effects models is slightly nearer to the nominal value its fixed-effects counterpart (Fig. 5a). Additionally, we found that the number of levels most strongly influences the type I error rate for mixed-as well as fixed-effects model (Fig. 5c). With five or more levels, however, the influence is almost zero. Differences between the mixed- and fixed-effects models arose for the variance and the total number of observations. Here, the mixed-effects model was less influenced by a small random effects’ variance and a low number of total observations than the fixed-effects model (Fig. 5 b, d). Balance did not influence both models (Fig. 5e).

**Figure 5:**
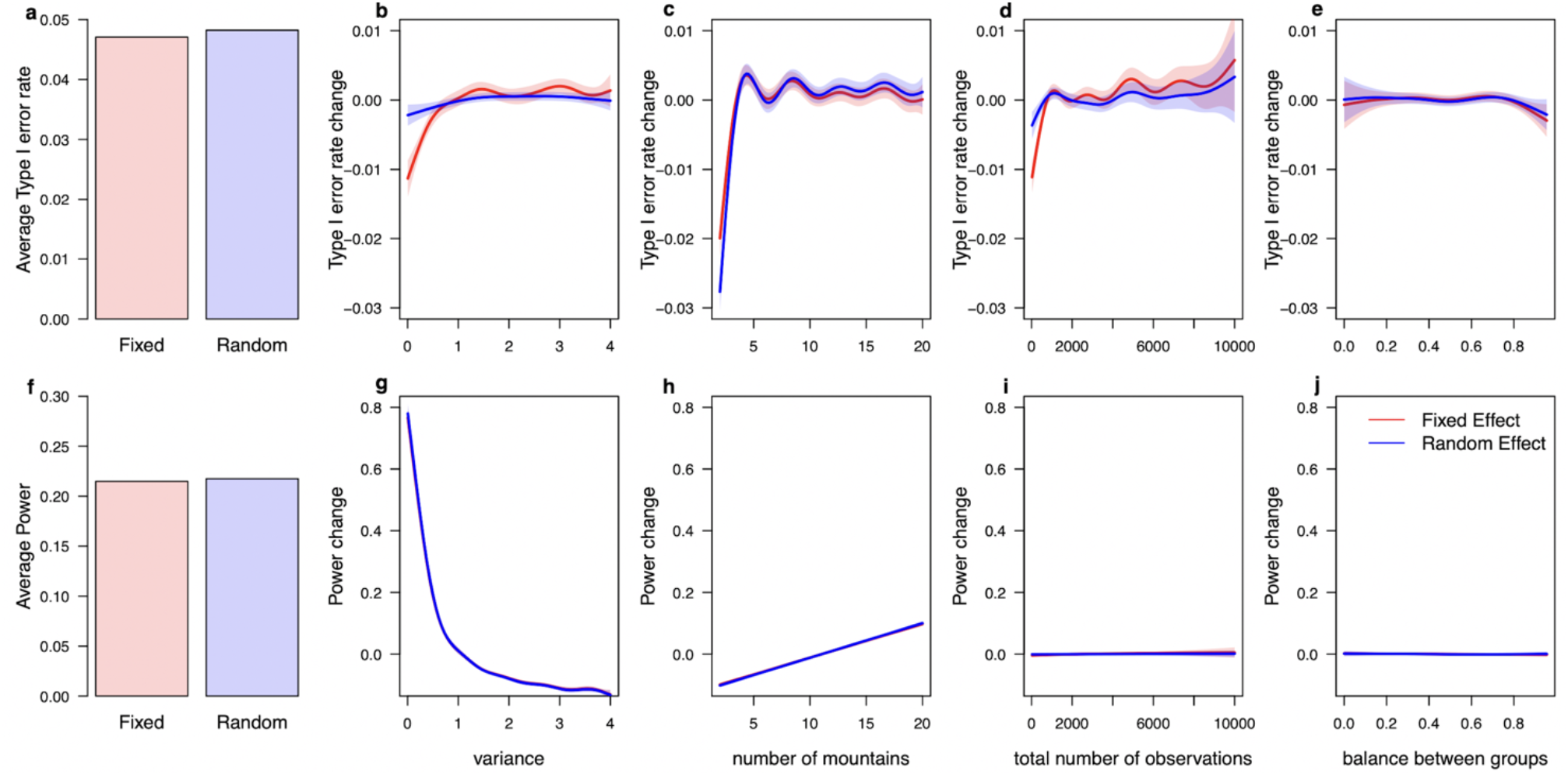
Comparing the influence of study design factors on the type I error rate (b - e) and power (g - j) of linear mixed- (blue lines) and fixed-effects models (red lines) with their respective average values (a, f). We found that the variance of the random effects and the number of levels (*number of mountains*) are the most important values to get correct statistical properties and improve the chance to find a significant effect. For this analysis, we used the plant height example (Hypothesis 2, Box 1, normal distribution models) for Scenario B (random intercept and random slope). Results for mixed-effects models are only from datasets in which mixed-effects models converged without presenting singular fit problems, while results for fixed-effects model are from all datasets.

For power we found no difference between fixed- and mixed-effects model (Fig. 5 f-j). For both models, an increase in variance decreased the power, while increasing the number of levels increased the power (Fig. 5 g, i). The total number of observations and the balance between groups had less influence (Fig. 5 h, j).

## Discussion

Ecological data collections or experiments often produce data with grouping/blocking structures and mixed-effects models can account for these dependencies. When such data, however, has a small number of levels, should the analysists stick to the mixed-effects model or fall back to a fixed-effects model? How does this decision influence the statistical properties of the population effect? Here, we showed with simulations that mixed-effects models with small number of levels in the grouping variable are technically more robust than previously assumed (Fig. 2) and that the decision between random and fixed effect matters most when the ecological effect differs among levels (Fig. 2).

When the ecological effect of interest is the same for each level of the grouping variable (scenario A, varying intercept, random intercept model), almost all models independent of the number of levels presented the same statistical properties for the ecological effect (temperature, slope) (Fig. 1, see also Gomes 2021). The only exception was the too complex model that presented too low type I errors (close to zero, Fig. 1) and lower power. We speculate this is caused by the higher number of parameters that need to be estimated by the model and that the model was unable to correctly estimate the random slopes to zero. Notably, for scenario A, the too simple model omitting the grouping variable presented correct statistical properties (Fig. 1). However, this is illusive because power decreased strongly with increasing variance in the random effect (Fig. S12) confirming the importance of including grouping variables to correctly partition the variance among the different predictors to get well calibrated statistical properties (Gelman 2005; Gelman & Hill 2007; Bell *et al*. 2019). Also including the grouping variable is mandatory if one is interested in the average intercept, otherwise it would cause inflated type I error rates (see Fig. S9, S11; see the following section).

When the ecological effect of interest differs for each level of the grouping variable (scenario B; varying intercept and slope; random intercept and random slope model), the model choice influenced the statistical properties more strongly. The mixed-effects model had a better type I error than the fixed-effects model, especially for a larger number of mountains (Fig. 2). Power was comparable for LMMs, but for GLMMs the mixed-effects model had higher power which may be linked to the distributional assumption of the random effects (see Gelman & Hill 2007): the regularization of the random effects, which stems from their distributional assumption, reduces the “effective” number of parameters as individual estimates are constrained to the mean. In summary, mixed-effects models have better type I error rates and higher power than fixed-effects models for linear and generalized models.

The different patterns between the linear and generalized models are mainly due to our bootstrapping approach to calculate the average effect and its standard error. For generalized models, the bootstrapping approach showed on average a slightly deflationary distribution of p-values (Fig. 18) leading to more conservative type I error rates. However, the properties of alternatives such as the zero-sum contrasts were even worse for different simulation scenarios (Fig. 18, Fig. S20).

Too complex mixed-effects models presented in both scenarios slightly lower type I error and power compared to the correctly parameterized mixed-effects model (Fig. 1, 2). This trade-off between type I error and power is in line with Matuschek *et al*. (2017) for different model complexities. Overall, the more complex (higher parametrized) models are more conservative but have less power than the simplified models. We think these more conservative estimates are preferable over anti-conservative estimates because researchers unfortunately tend to try a variety of analyses and only report significant ones (Simmons *et al*. 2011) and more conservative type I error rates counteract this procedure.

However, dropping the correlation structure between random effects should be still carefully considered. Correlations between the random effects are very common in empirical data and the true change in type I error rate might be higher than shown here because we assumed no correlation in our simulations. Group-mean centering of the ecological effect may mitigate the requirement of assuming a correlation, but it also changes the interpretation of the model.

In scenario B, too simple models exhibited inflated type I errors (in line with Schielzeth & Forstmeier 2009; Barr *et al*. 2013; Bell *et al*. 2019), but very high power (Fig. 2). We speculate that additional variance coming from the difference between levels in the grouping variable, which is not accounted, is attributed to the ecological effect and causes overconfident estimates.

### Variances of random effects and singular fits

Singular fits occurred more often for small number of levels, creating a point mass at zero in the distribution of the variance estimates (Fig. 3, Table S1), because a singular fit in our simulations corresponded to a zero-variance estimate as we excluded the other possibility (we set the correlations between random slope and intercept to zero, see *Methods*). The resulting distribution consisted of a towards higher values biased chi-squared distribution and the point mass at zero as expected (see Stram & Lee 1994). For non-singular and singular fits combined, the variance estimates of the random effects were biased and imprecise with a small number of levels, but the bias decreased with the number of levels towards zero (McNeish 2017). Removing the singular fits led to even more bias in the variance estimates (Fig. 3c, d).

The biased variance estimates are caused by ensuring positive variances in the optimization routines (Bates *et al*. 2015; Brooks *et al*. 2017). In case of a singular fit, the correctly specified mixed-effects model had similar statistical properties as a fixed-effect model dropping the grouping variable (Fig. 4): no difference between the levels, which corresponds to a fixed-effect model without the grouping variable. However, the models still differed in their number of parameters (and degrees of freedom) which might explain the slight differences in statistical properties (Fig. 4).When switching to fixed-effects models for singular fits in the random effect, the type I error rate and power were similar to the random effect model with non-singular fits (Fig. 4).

### Connection to study design

Earlier studies reported mixed recommendations about important study design factors. While some studies only stressed the importance of the total number of observations (Martin *et al*. 2011; Pol 2012), we found, in accordance with Aarts *et al. (*2014), that the number of levels and the variance between levels have a strong influence on type I error rates and power. Due to our simulation design, which automatically increases the number of observations when increasing the number of levels, we however, cannot perfectly separate the effects of number of observations and levels from each other.

The influence of the variance on the statistical properties is mixed. On the one hand, increasing the variance had a positive effect on the type I error for both models but the fixed-effects model was more strongly affected (Fig. 5). The different distributional assumptions might explain this different behavior: the mixed-effects model assumes the levels to be normally distributed and estimates flexibly the variance of the levels, whereas the fixed-effects model makes implicitly the assumption of an infinite variance. On the other hand, increasing the variance over a certain value (Fig. 5g) decreased the power of both models because more variance is explained by the difference between levels, and this increases the uncertainty of the slope effect estimate.

Given the strong influence of the number of mountains on type I error rates, we encourage to design a study with at least 8 levels because with more than 8 levels, the type I error rate was approximately not affected by the number of mountains (Fig. 5c). In our scenarios, the influence of the unbalanced number of observations between levels was small (Fig. 5) suggesting that unbalanced designs are not the main concern for mixed-effects models given their robustness to unbalanced data (Swallow & Monahan 1984; Pinheiro & Bates 1995; Schielzeth et al. 2020) as long as one is not interested in treatment but population effects. Moreover, the impact of study design on type I error and power stresses the importance of pre-experiments and power analyses (e.g. Johnson *et al*. 2015; Green & MacLeod 2016; Brysbaert & Stevens 2018) to maximize the meaningfulness and efficiency of a study.

### Practical suggestion

Before giving practical advice, we must recall the exact situation in which this manuscript acts. We assume that a researcher is interested in a population effect, and they have already decided to use a mixed-effects model (broad-sense analysis, not interested in the individual levels effects), but now faces a low number of levels. Recommendations only apply to such situations.

We found no harsh threshold for the minimum number of levels in a grouping variable necessary to model it as a random effect (Fig. 2), but rather found that a singular fit in the mixed-effects model indicates switching to the fixed-effects model is beneficial (more conservative type I error rates, Fig. 2). Acknowledging that most singular fits occur with a small number of levels (Table S1), this might also explain the common rule of thumb to do not fit a grouping variable as random effect if it has fewer than 5 levels (Gelman & Hill 2007; Bolker et al. 2009; Bolker 2015). Apart from the results of our study, checking the data for possible issues causing zero variance estimates is desired. Additionally, one should investigate whether a zero-variance estimate is ecologically reasonable.

We therefore recommend starting with the mixed-effects model, regardless of the number of levels, and switching to a fixed-effects model only in case of a singular fit (more conservative type I error rate). When the ecological effect differs among levels, we recommend starting with correlated random slope and intercept (following Barr et al. 2013). When obtaining a singular fit, switch to uncorrelated random effects (following Matuschek *et al*. 2017) and in case of another singular fit, switch to a fixed-effects model. These recommendations are summarized in Fig. 6.

**Figure 6:**
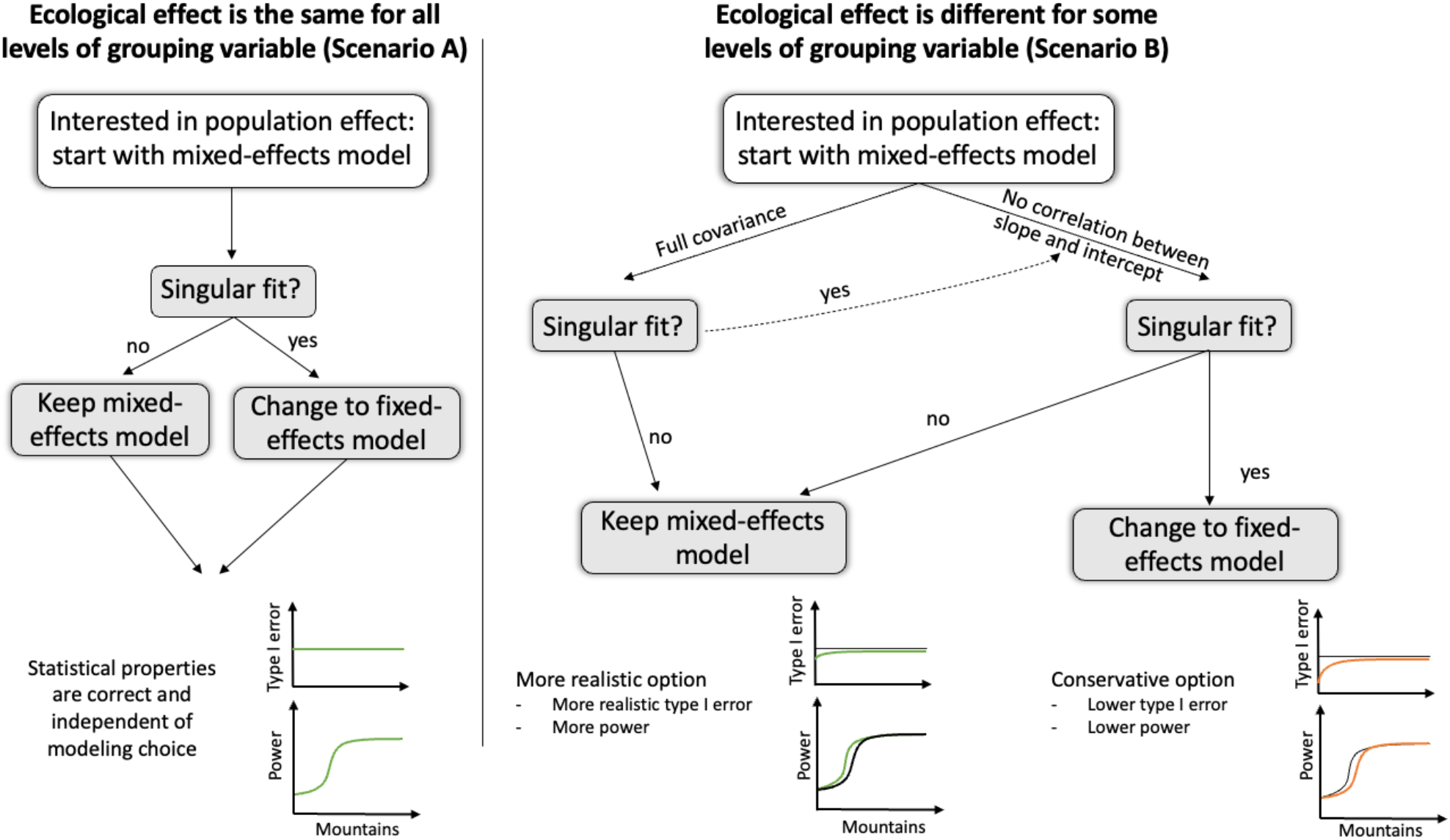
Consequences and recommendations for mixed-effects models with a small number of levels in the random effect. When the ecological effect does not differ between different levels of the grouping variable (left side) all modeling options, which include the grouping variable, lead to the same results and thus only a singular fit requires a change to a fixed-effects model. If the ecological effect differs among levels (middle to right side), starting with the mixed-effects model and only changing to the fixed-effects model in case of a singular fit is recommended.

Our previous recommendations assumed that we know whether the ecological effect differs across levels. With empirical data, however, this is usually unknown. Here, a too complex model leads to too low type I error rates and less power, while a too simple model has inflated type I error rates. Thus, to prevent being overly confident, it is essential to include random slopes if they are present (Schielzeth & Forstmeier 2009). But because of the power issue, we doubt that its generally advisable to always start with the maximally complex random effect structure (as suggested in Bell *et al*. 2019), and it requires a decision criterion for the complexity of the random effect structure.

Unfortunately, model selection on random effects is complicated, because the exact degrees of freedom, that are used by a random effect, are unclear (Kuznetsova *et al*. 2017). Approximations exist and for low number of levels a Kenward-Rogers correction is preferred (McNeish 2017). Although this prevents a naïve use of AIC or likelihood ratio tests (LRT), other methods such as simulated (restricted) LRTs (Wiencierz *et al*. 2011) can be used to decide if adding a random slope is justified. Thus, we recommend starting with the simpler structure (typically random intercepts) and then use residual checks (e.g. Hartig 2019) or appropriate model selection criteria (e.g. Matuschek *et al*. 2017) to decide if a random slope should be added.

Another, somewhat perpendicular option is to modify the distributional assumption about the random effects (normal distribution), which is mainly due to mathematical convenience (Beck & Katz 2007) and corresponds to a L2 (RIDGE) regularization. A L2 regularization biases random effect estimates towards zero but is unable to produce exactly zero estimates (Zou & Hastie 2005). In contrast, L1 (LASSO) regularization, which corresponds to a Laplace distribution (Park & Casella 2008; Hans 2009; Tibshirani 2011), is able to estimate zero random effect estimates, when there is no effect. Because mixed-effects models are relatively robust to distributional assumptions (Bell *et al*. 2019; Schielzeth *et al*. 2020), switching distributions to LASSO or a combination of L1 and L2 regularization (elastic net, see Zou & Hastie 2005) could potentially allow starting with the maximal complex random effect structure (see also McNeish & Bauer 2020). Such options should be investigated in further studies.

## Conclusion

In conclusion, we showed that mixed-effects models are more robust than previously thought, despite the biased variance estimates for low number of levels in the grouping variable. We found that the statistical properties of the population effect are robust against the model choice when the ecological effect is the same among the levels of the grouping variable, however, the model matters when the ecological effect differs among levels. When in doubt about the data-generating process, we encourage starting with a simplified model (random intercept only) and consult model diagnostics and simulated LRTs to check for evidence of random slope effects. When finding evidence for random slopes in these tests, we recommend starting with the mixed-effects model and switching only to a fixed-effects model in case of a singular fit problem. With this work, we provide a practical guideline, which helps ecologists in the study design, the data analysis and thus making ecological inference more informative and robust.

## Data availability statement

No empirical data was used in this study. Code to run and analyze the experiments can be found at https://github.com/JohannesOberpriller/RandomEffect_Groups.

## Author Statement

MP, JO and MSL designed the study. MP and JO ran the simulations, analyzed the results and wrote a first draft. All authors contributed equally to revising the manuscript and interpreting and discussing results.

## Acknowledgement

The idea of the manuscript originated from a discussion in the Theoretical Ecology seminar and was further developed in the Coding Club at University of Regensburg. We thank Rainer Spang, Carsten Dormann, Magdalena Mair, Björn Reineking, Sean McMahon, Andreas Ettner and Florian Hartig for comments and discussions on earlier versions of the manuscript. We also thank two anonymous reviewers for their valuable comments and suggestions. JO was funded by the Bavarian Ministry of Science and the Arts in the context of Bavarian Climate Research Network (bayklif). MSL was funded by the Smithsonian Predoctoral Fellowship.

