## Supporting Information S1 for "Fixed or random? On the reliability of mixed-effects models for a small number of levels in grouping variables"

### 1. Mixed-effect model implementations in R, default settings and convergence issues

The most used packages in R to fit mixed-effect models are *lme4* (Bates et al., 2015) and *glmmTMB* (Brooks et al., 2017). These packages differ in their optimization routines and the calculation of p-values for linear mixed-effect models (LMMs). While *lme4* uses standard optimizers, *glmmTMB* relies on automatic differentiation implemented in *TMB* package (Kristensen et al., 2016). Another difference is that *glmmTMB* offers to fit linear and generalized linear models with both the maximum likelihood (MLE) and the restricted maximum likelihood estimation (REML), while *lme4* offers only REML for LMMs but not for GLMMs. By default, *lme4* uses restricted maximum likelihood (REML) for LMMs (*lmer* function) and unrestricted maximum likelihood (MLE) for GLMMs (*glmer* function), while *glmmTMB* uses MLE by default for any kind of data. Due to these different optimization routines and default options, *glmmTMB* and *lme4* results could be slightly different, which we analyze in the following.

#### 1.1 Variance estimates and singular fits

With *glmmTMB* using REML the estimates of variances consist of a point mass at zero and a Chi-squared distribution (Fig. S1). When excluding simulations which presented variance estimates smaller than $10^{-3}$, the point mass at zero vanishes (Fig. S2). These estimates correspond to singular fits obtained with *lme4*.

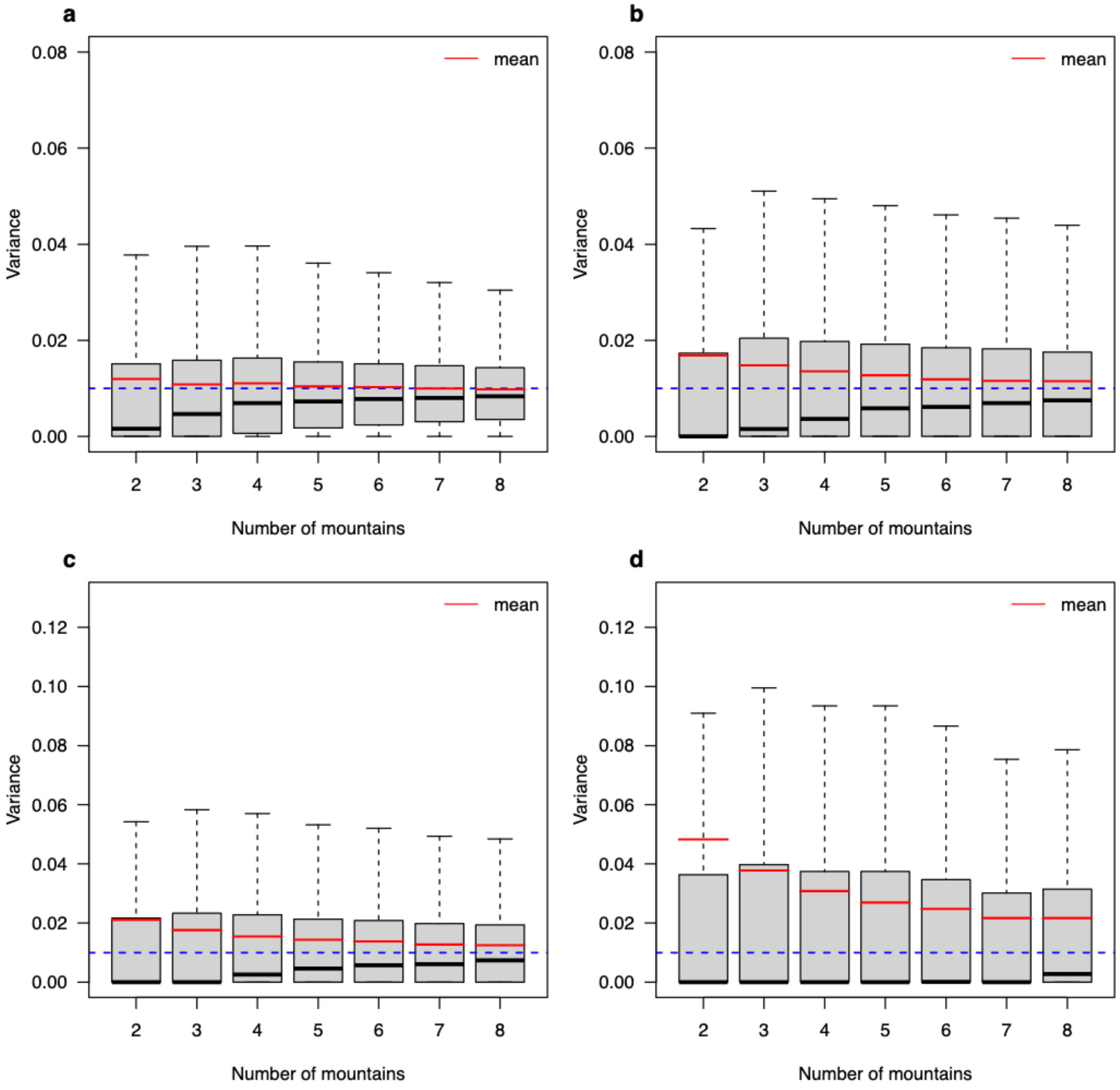


**Figure S1**: Variance estimates of the random intercepts (a, b) and random slopes (b, d) for linear mixed-effect models (a, b) and generalized linear mixed-effect models (c,d) from the correctly specified mixed-effect model

(Table 1. Eq. 10) in Scenario B, fitted with *glmmTMB* package using REML to simulated data with 2-8 mountains. For each scenario, 5,000 simulations and models were tested. The grey lines represent the true variance used in the simulation (0.01).

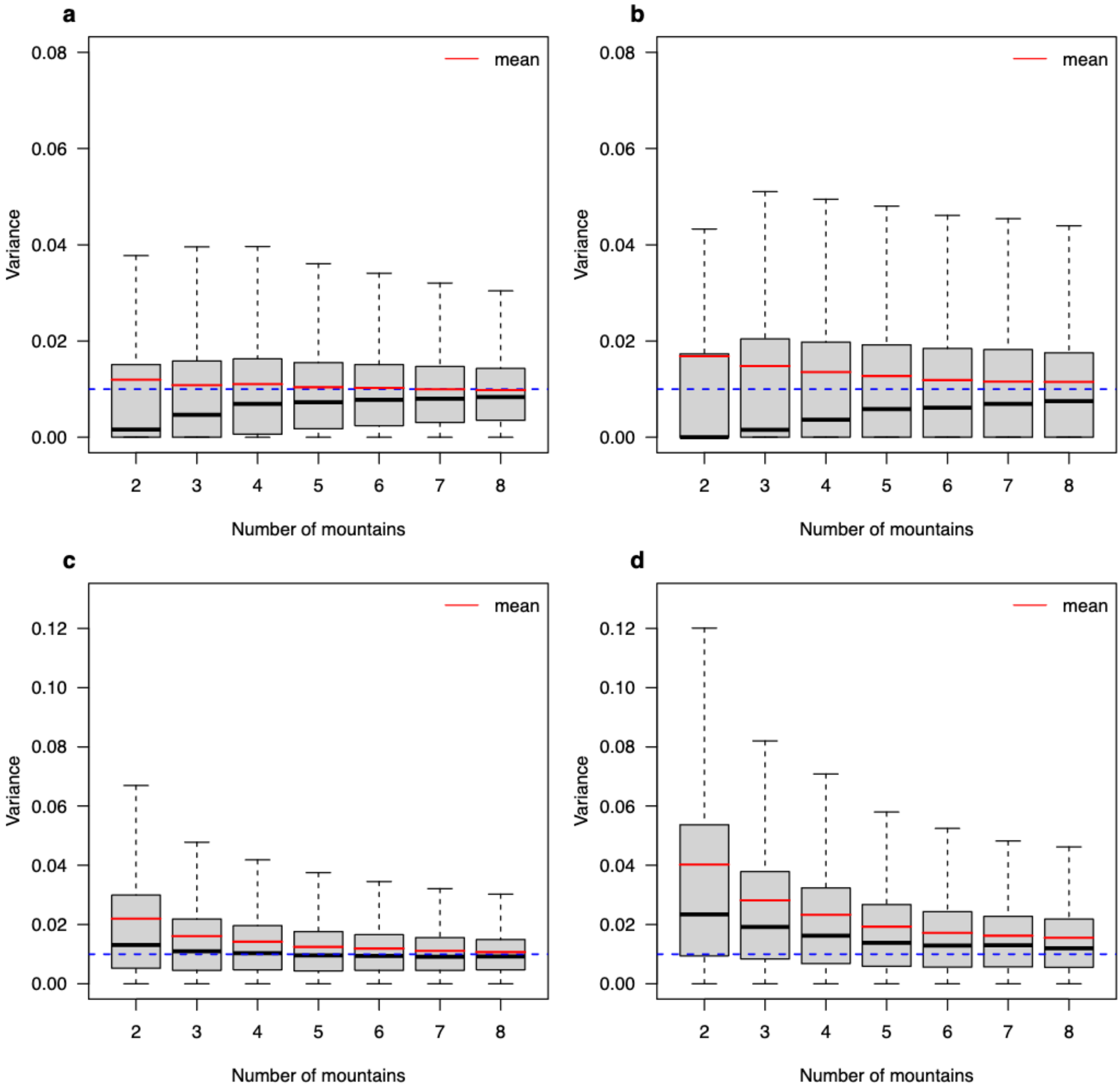


**Figure S2**: Variance estimates of the random slopes and random intercepts (model 4) for linear mixed-effect models (LMM) fitted with *glmmTMB* using REML to simulated data with 2-8 mountains. (a) and (b) show the results for all models (with and without singular fits), (c) and (d) show the results for the models without what we considered as singular fits (variance lower than $10^{-6}$). For each scenario, 5,000 simulations and models were tested. The grey line represents the true variance used in the simulation (0.01).

In Table S1, we compare the distributions of the variances of the random effects (random intercept and slope) of the correctly specified model in scenario B between REML and MLE using the package *glmmTMB,* due to its flexibility in fitting models with both algorithms.

**Table S1:** Proportion of models ran in *lme4* that presented singular fit convergence problem when using maximum likelihood (MLE) and restricted maximum likelihood (REML) fitting algorithms. Notice that for GLMMs in *lme4*, REML is not implemented.

| | **LMM** | | **GLMM** |
|---|---|---|---|
| **Number of groups** | **REML** | **MLE** | **MLE** |
| 2 | 77% | 92% | 95% |
| 3 | 64% | 80% | 89% |
| 4 | 55% | 70% | 84% |
| 5 | 45% | 61% | 81% |
| 6 | 40% | 54% | 78% |
| 7 | 36% | 48% | 76% |
| 8 | 32% | 43% | 72% |

Additional to the number of singular fits a direct comparison of REML and MLE with respect to their estimates of the variances is necessary to compare their performance. Irrespective of the specific package (*lme4* Fig. S3 A, B or *glmmTMB* Fig S3 c, d) using MLE for linear mixed-effect models lead to estimates, which are stronger biased towards zero than REML estimates. The same applies for generalized mixed-effect models (Fig. S4).

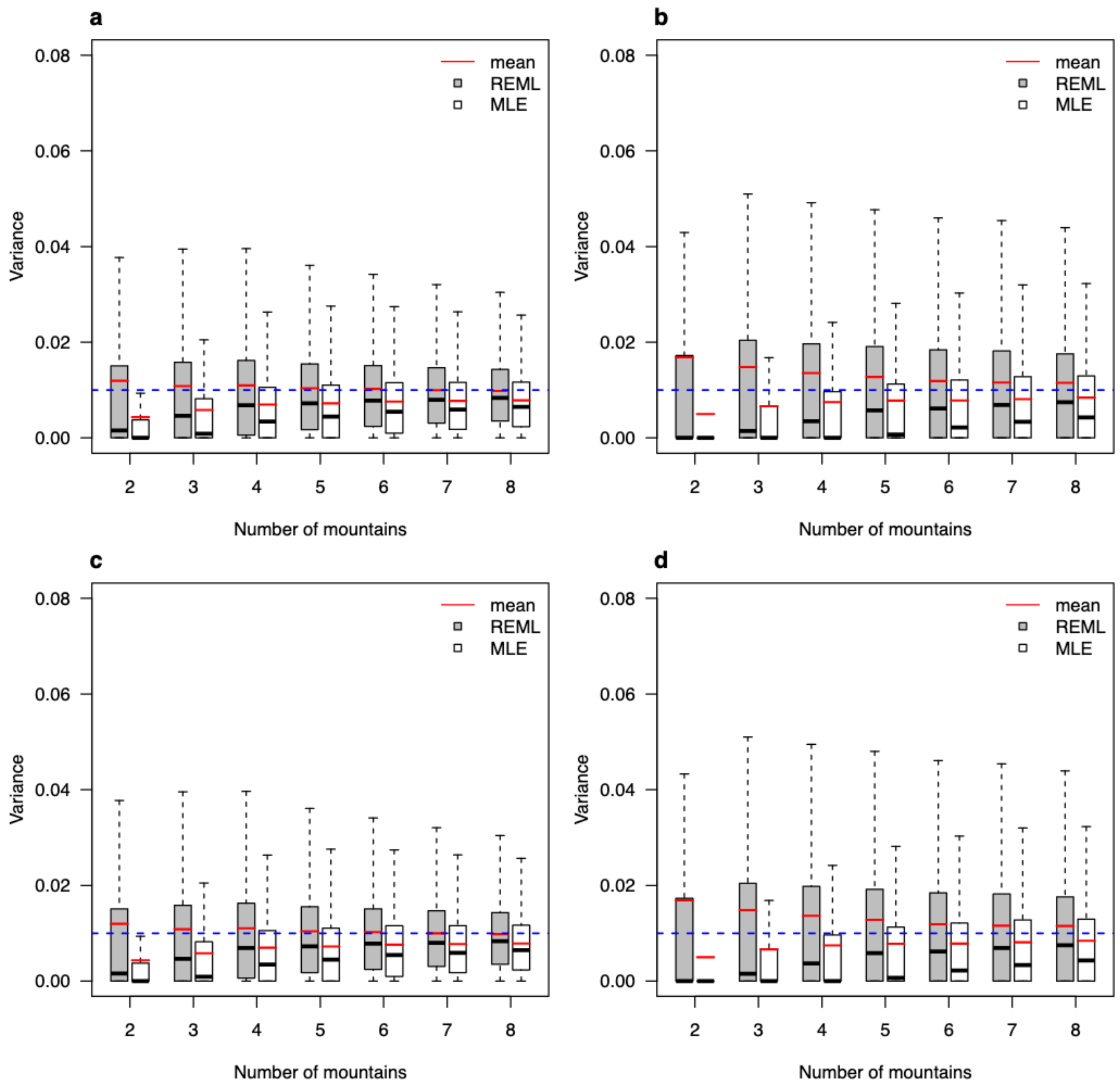


**Figure S3**: Variance estimates of the random intercepts (a, c) and random slopes (b, d) for **linear mixed-effect models** (LMM) fitted to simulated data with 2-8 numbers of artificial mountain ranges. For each scenario, 5,000 simulations and models were tested. The blue line represents the true variance used in the simulation (0.01). (a) and (b) show the results for linear mixed-effect models fitted with the *lme4* R package, (d) and (d) show the results for linear mixed effects models fitted with the *glmmTMB* package. The grey boxes show results for the models fitted by restricted maximum likelihood estimation (REML) and the white boxes shows results for the models fitted by maximum likelihood estimation (MLE).

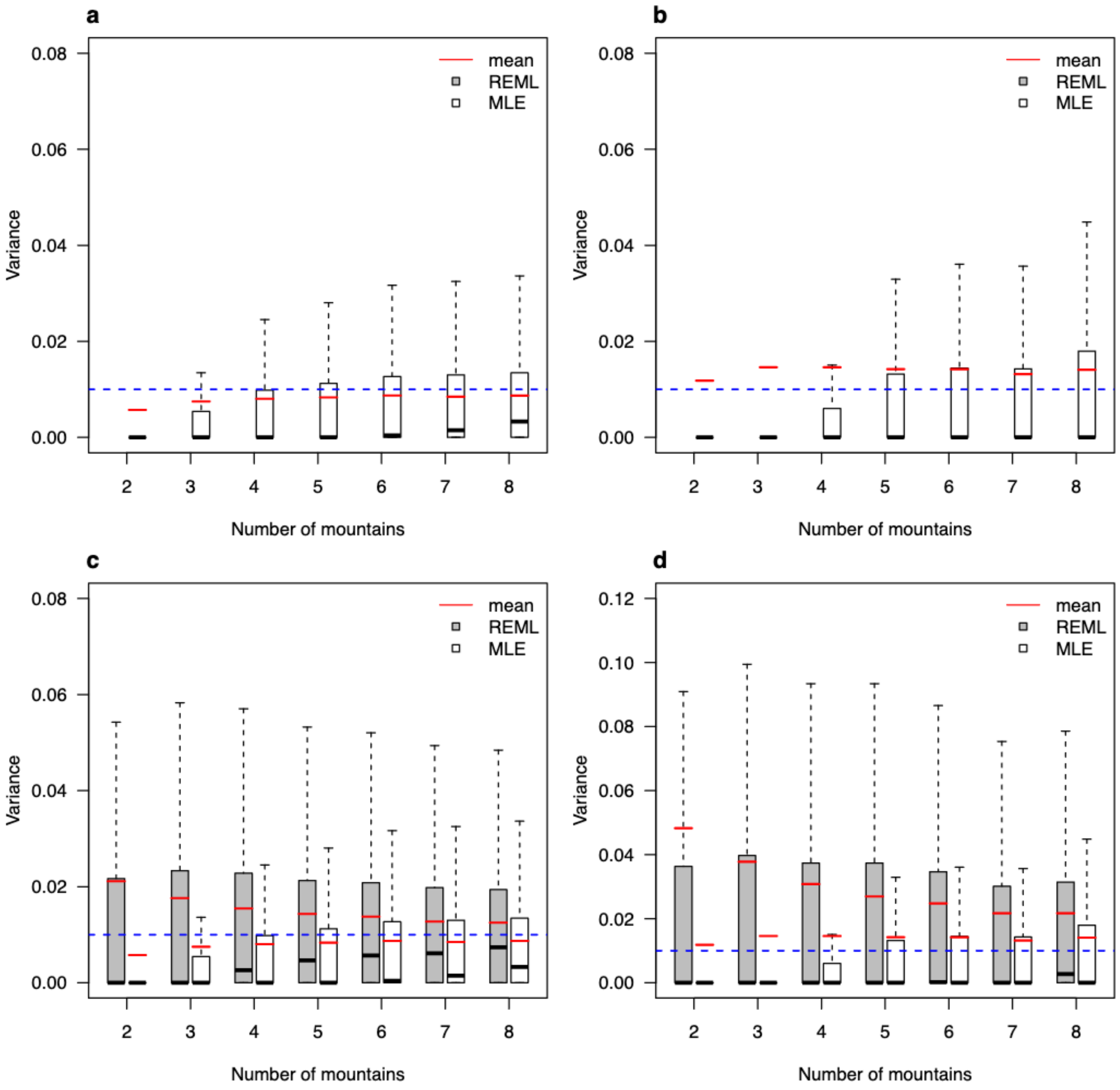


**Figure S4**: Variance estimates of the random intercepts (A, C) and random slopes (B, D) for **generalized linear mixed-effect models** (GLMM) and linear regression models fitted to simulated data with 2-8 numbers of artificial mountain ranges. For each scenario, 5,000 simulations and models were tested. The blue line represents the true variance used in the simulation (0.01). (a) and (b) show the results for GLMMs fitted by the lme4 R package, (c) and (d) show the results for GLMMs fitted by the *glmmTMB* package. The grey boxes represent the results for models fitted by restricted maximum likelihood estimation (REML) and the white boxes shows the results for models fitted by maximum likelihood estimation (MLE).

Additional to the number of singular fits, a direct comparison of imbalanced and balanced data both fitted with REML with respect to the estimates of the variances is necessary to compare the influence of unbalance between groups. Irrespective of the specific package (*lme4* Fig. S5a,b or *glmmTMB* Fig S5c,d) estimates for balanced and unbalanced data do not differ. For generalized mixed-effect models the same applies (Fig. S6).

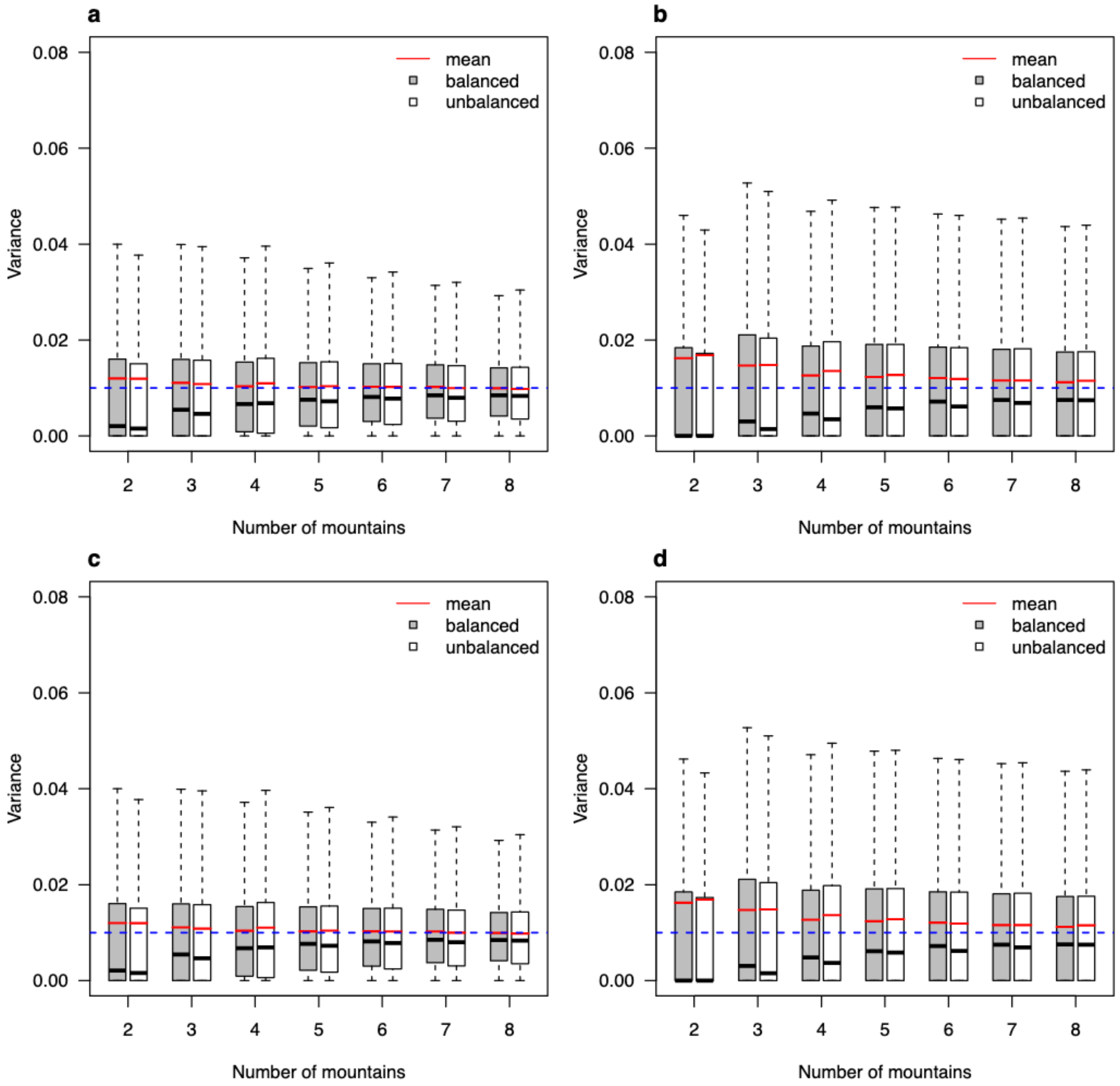

**Figure S5**: Variance estimates of the random intercepts (a, c) and random slopes (b, d) for **linear mixed-effect models** (LMM) fitted to simulated data with 2-8 numbers of artificial mountain ranges using REML. For each scenario, 5,000 simulations and models were tested. The blue line represents the true variance used in the simulation (0.01). (a) and (b) show the results for linear mixed-effect models fitted with the *lme4* R package, (c) and (d) show the results for LMMs fitted with the *glmmTMB* package. The grey boxes show the results for the models with unbalanced data (number of observation) among mountains and the white boxes shows results for the models fitted with balanced data among mountains.

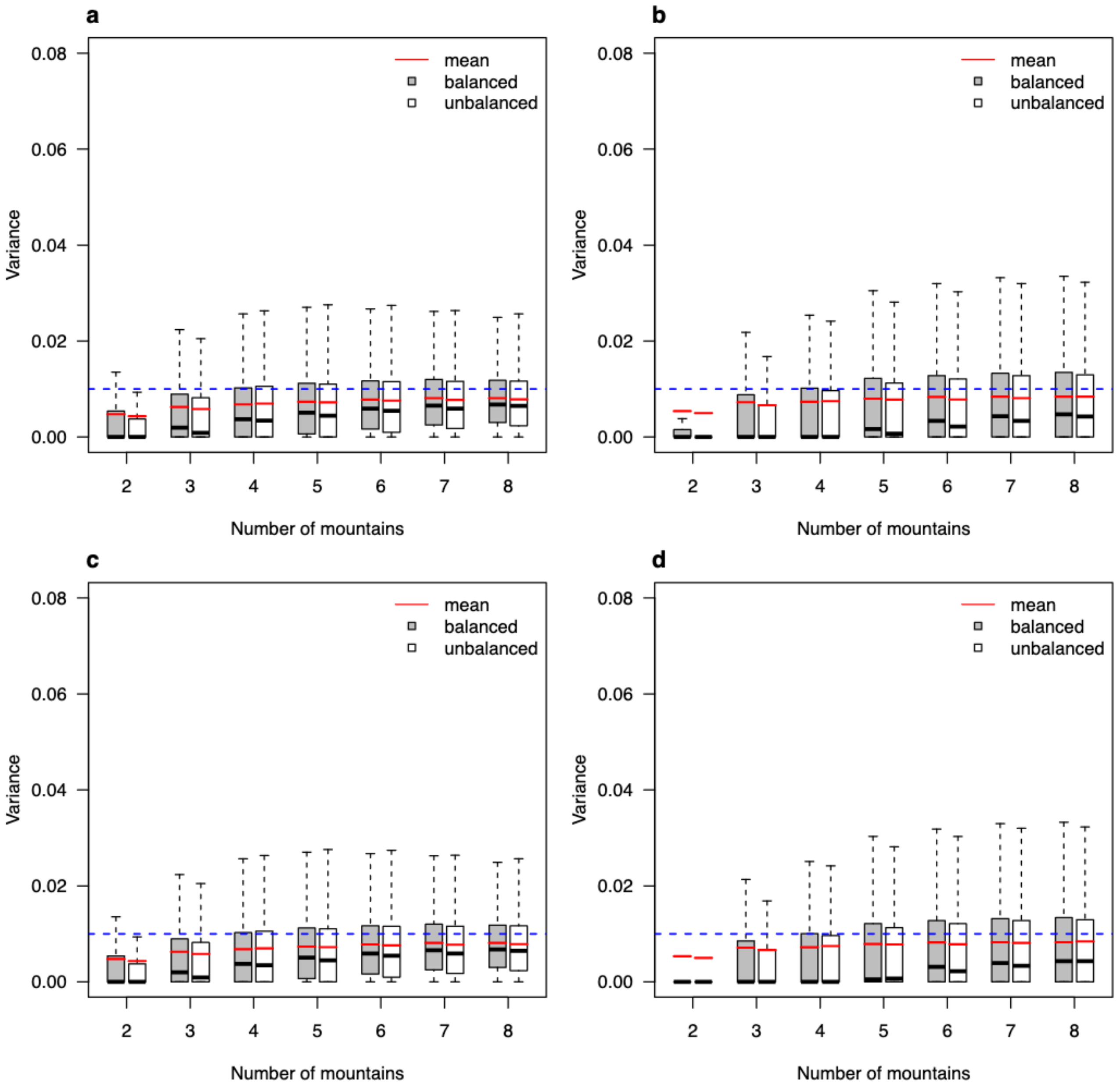


**Figure S6**: Variance estimates of the random intercepts (a, c) and random slopes (b, d) for **linear mixed-effect models** (LMM) fitted to simulated data with 2-8 numbers of artificial mountain ranges using MLE. For each scenario, 5,000 simulations and models were tested. The blue line represents the true variance used in the simulation (0.01). (a) and (b) show the results for linear mixed-effect models fitted with the *lme4* R package, (c) and (d) show the results for LMMs fitted with the *glmmTMB* package. The grey boxes show the results for the models with unbalanced data (number of observation) among mountains and the white boxes shows results for the models fitted with balanced data among mountains.

### 1.2. p-value calculations for mixed effect models

There is also a difference in the calculation of p-values between the two packages for linear mixed-effect models (LMMs). While *lme4* uses a Satterhwaite approximation to calculate degrees of freedom which then are fed into t-statistics, *glmmTMB* uses z-statistics and thus avoids the calculation of degrees of freedom. Z-statistics are the asymptotic limits of t-statistics when having infinite data, however, these two differ in the low data limit and p-values calculated using z-statistics are overconfident compared to using t-statistics. We see this for Hypothesis 2 in our Scenario A (Fig. S7a), for which t-statistics can be calculated analytically: t-statistics lead to values around the nominal type I error rates, while z-statistics lead to increased type I error rates. This also translates into power, where using z-statistic causes higher, but probably too high power compared to t-statistic (Fig. S7a). For generalized linear mixed-effect models (GLMMs), both packages use z-values and, thus, present the same statistical properties (Fig. S7b). We speculate the reason why *glmmTMB* is using z-statistics also for LMMs is that t-statistics cannot be calculated analytically for GLMMs and must be approximated. However, when interpreting the results, it is important to keep in mind that z-statistics are only approximations in the low data limit. We believe that the power of GLMMs would be similar to power of LMMs when t-values would be used instead of z-values.

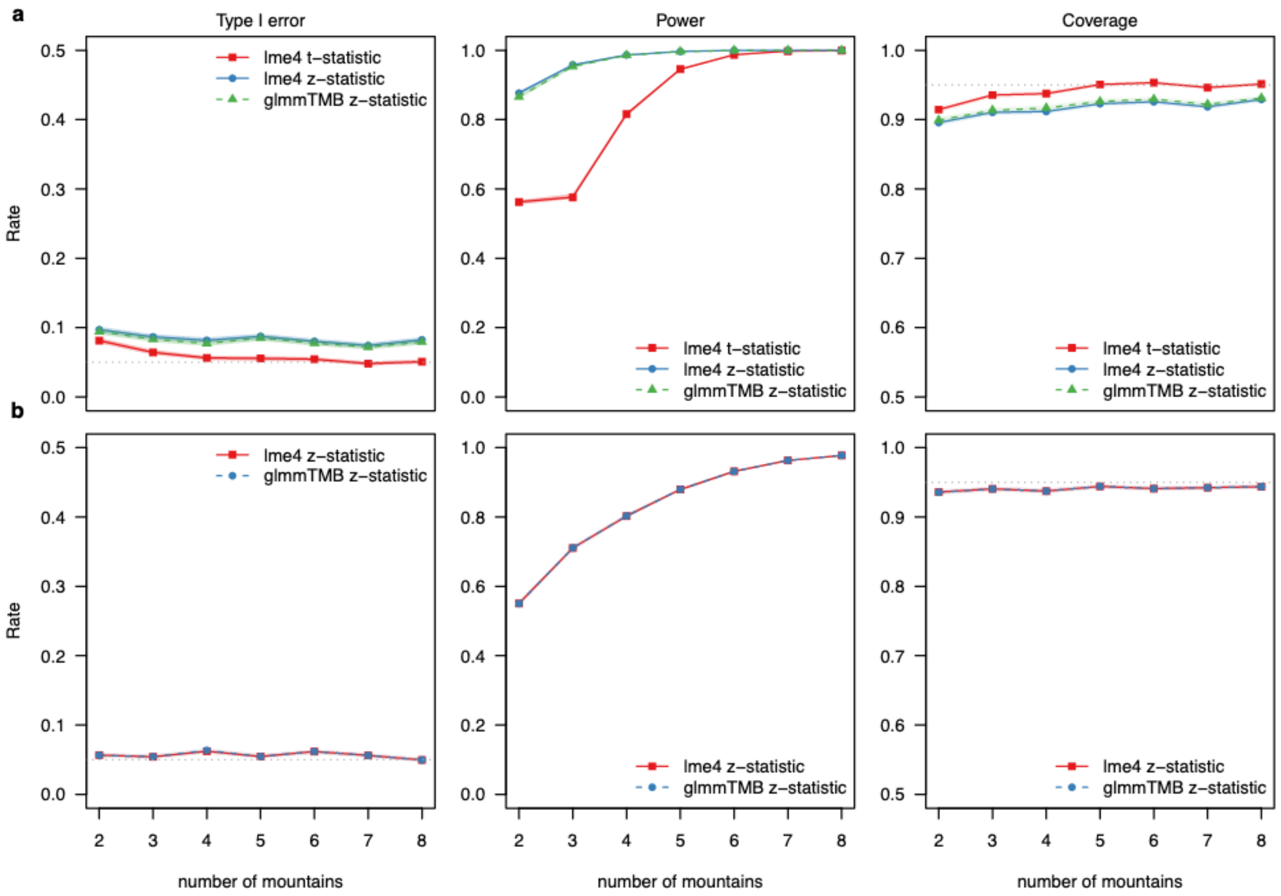


**Figure S7:** Comparing type I error rates, power, and coverage for linear mixed-effect models (a) and generalized linear mixed-effect models (b) between t- and z-statistics for *lme4* and *glmmTMB* packages. Data was simulated with 2-8 mountains for scenario A (random intercept only) with 50 (LMM) and 200 (GLMM) observations per mountain range. For each scenario, 5,000 simulations and models were tested.

### 2. Results with singular fits and results for intercepts

We calculated additionally the nominal type I error rate and power, i.e., with singular fits (Fig. S8, Fig. S10). As expected, the statistical properties are now approximatively around the nominal values (i.e., 5% for type I error rates).

Moreover, we calculated the statistical properties for the intercept estimates of Scenario A and B (Fig. S9, Fig. S11) and we found similar patterns as for the slope estimates (Fig. 1, Fig. 2).

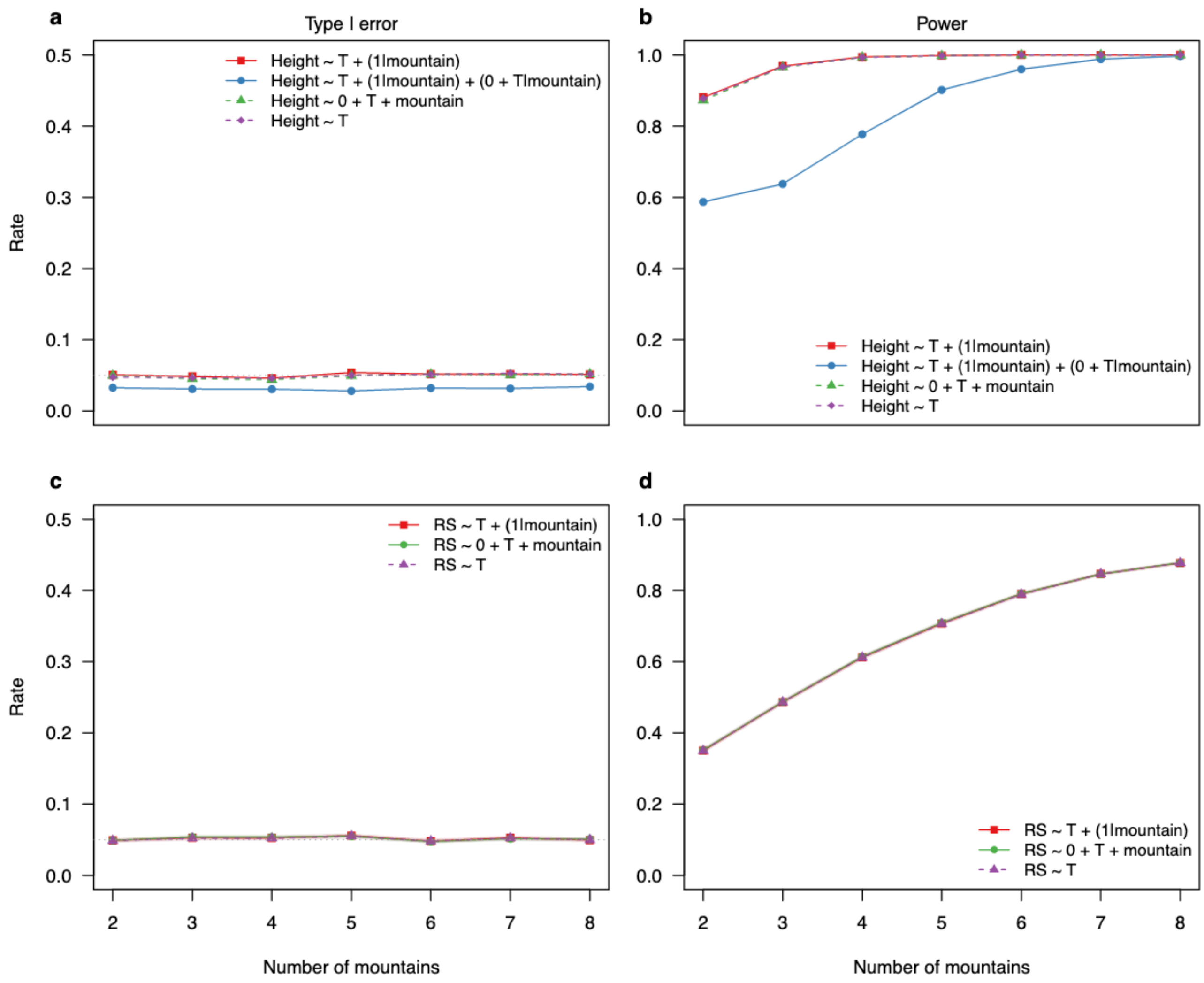


**Figure S8**: Type I error rates, and power for linear fixed and mixed effect models (a, b) and generalized linear fixed and mixed-effects models (c, d) with singular fits and non-singular fits combined in the results. Models were fitted to simulated data with 2-8 mountains (random intercept for each mountain - Scenario A) and 50 observations per mountain in *lme4* package. For each scenario, 5,000 simulations and models were tested.

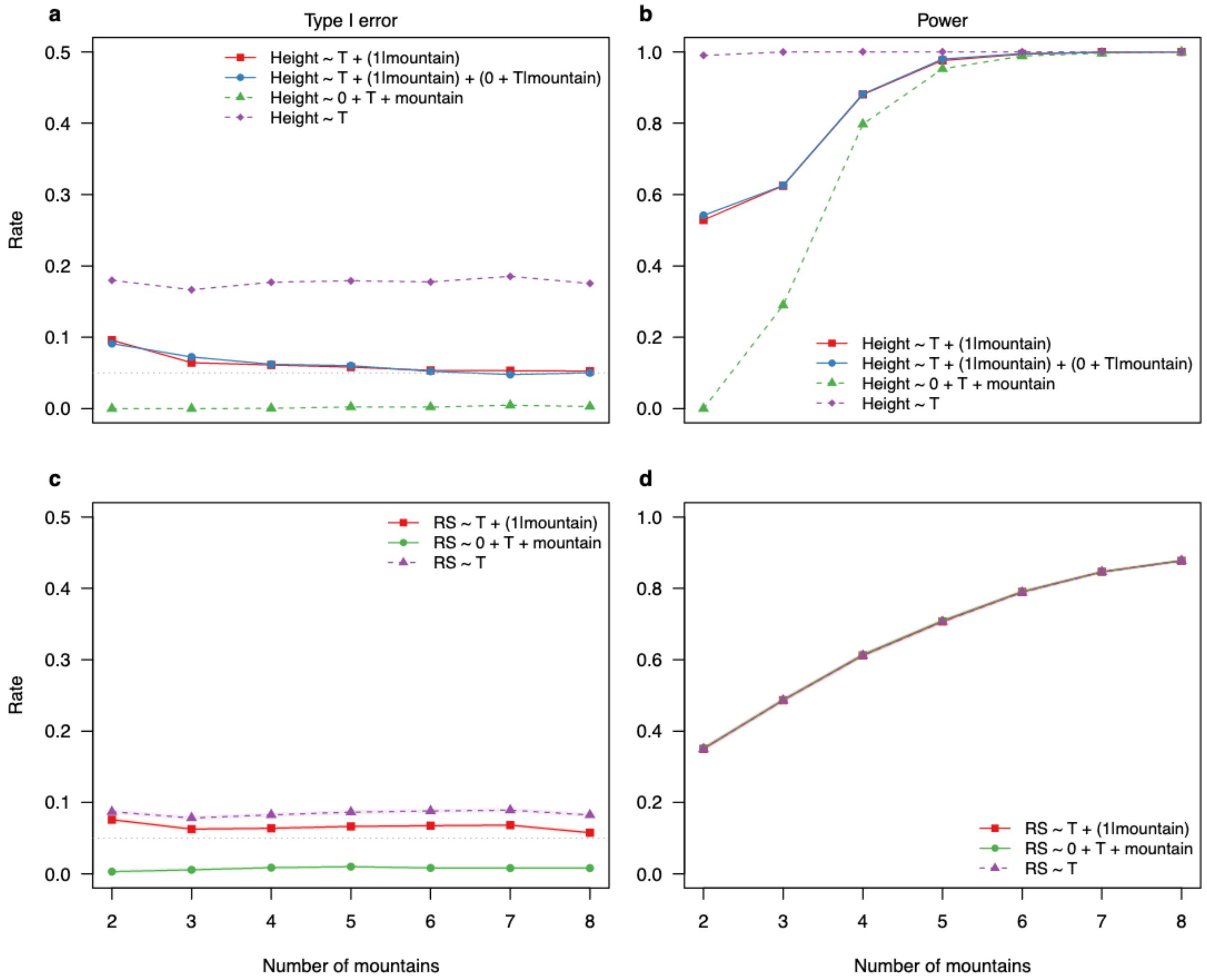


**Figure S9**: Type I error rates, and power for linear fixed and mixed effect models (a, b) and generalized linear fixed and mixed-effects models (c, d) for the average intercept, fitted to simulated data with 2-8 mountains (random intercept for each mountain - Scenario A, intercept) and 50 observations per mountain. For each scenario, 5,000 simulations and models were tested. These results are only for datasets in which mixed-effects models converged without presenting singular fit problems.

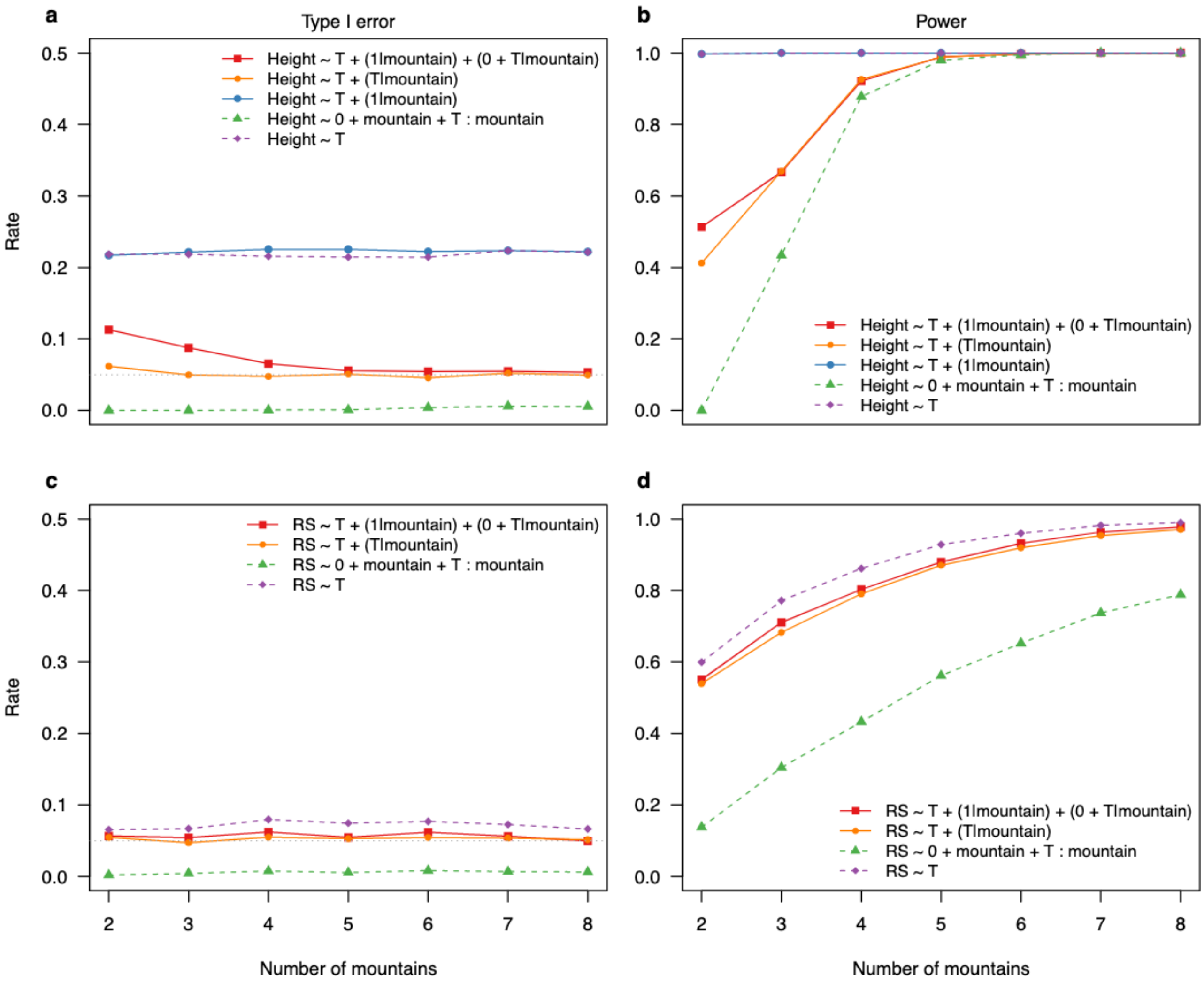


**Figure S10**: Type I error rates, and power for linear (mixed effect) models (a, b) and generalized linear (mixed-effects) models (c, d), fitted to simulated data with 2-8 mountains for scenario B (random intercept and random slope for each mountain range) with singular and non-singular fits combined in the results. For each scenario, 5,000 simulations and models were tested.

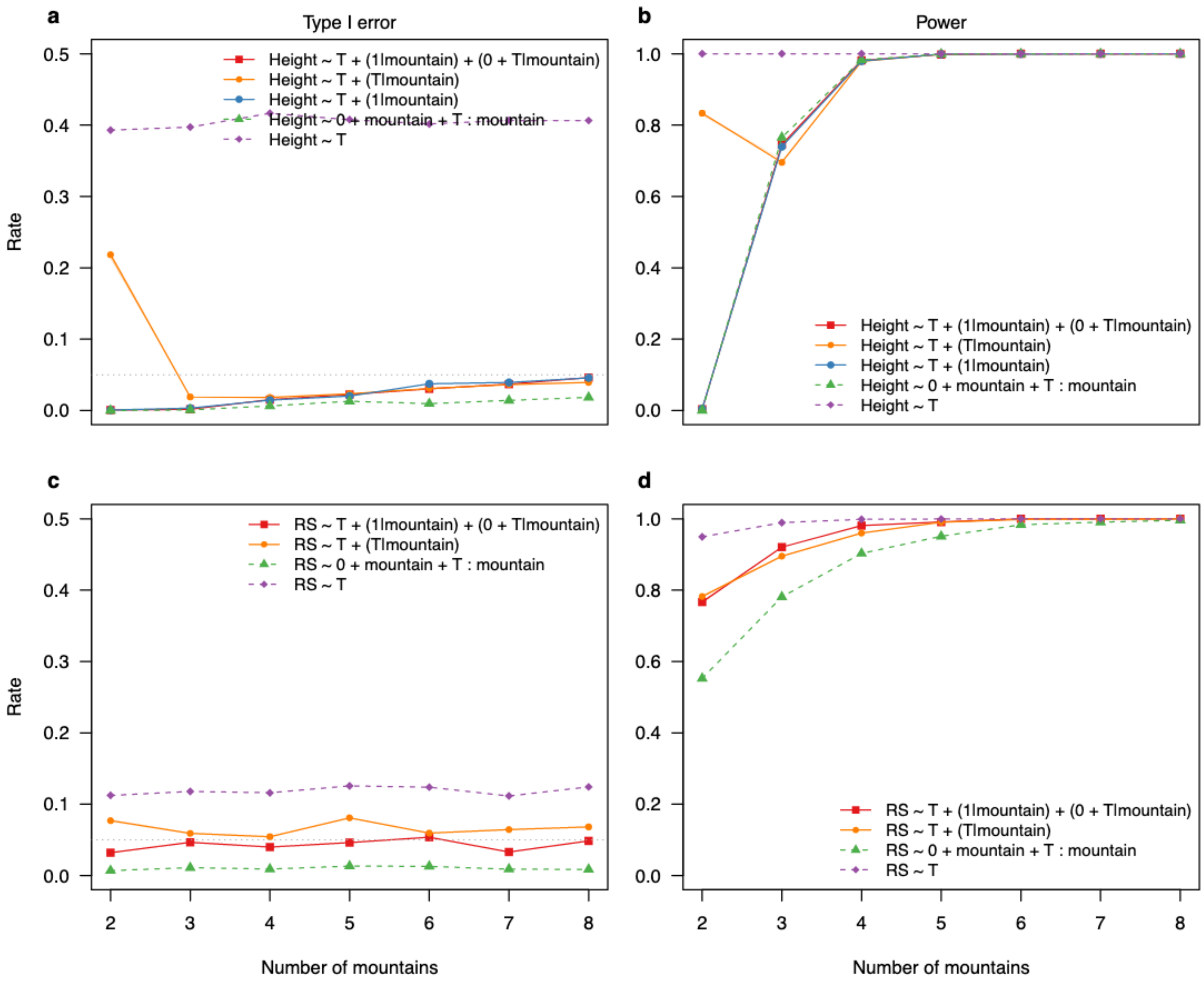


**Figure S11**: Type I error rates, and power for linear (mixed effect) models (a, b) and generalized linear (mixed-effects) models (c, d) (intercept), fitted to simulated data with 2-8 mountains for scenario B (random intercept and random slope for each mountain range). For each scenario, 5,000 simulations and models were tested.

### 3. Varying standard deviation for the random effects for linear models

Since we found that the variance of the grouping variable (mountains) influenced the statistical properties of the model (Fig. 5, main text), we ran additional data with 4 different standard deviations for the random effects in the data-generating model (0.01, 0.1, 0.5,

2.0) for the linear model example (Hypothesis 2, Box 1) and compared type I, power, and coverage and in both scenarios, A (Fig. S12) and B (Fig. S13).

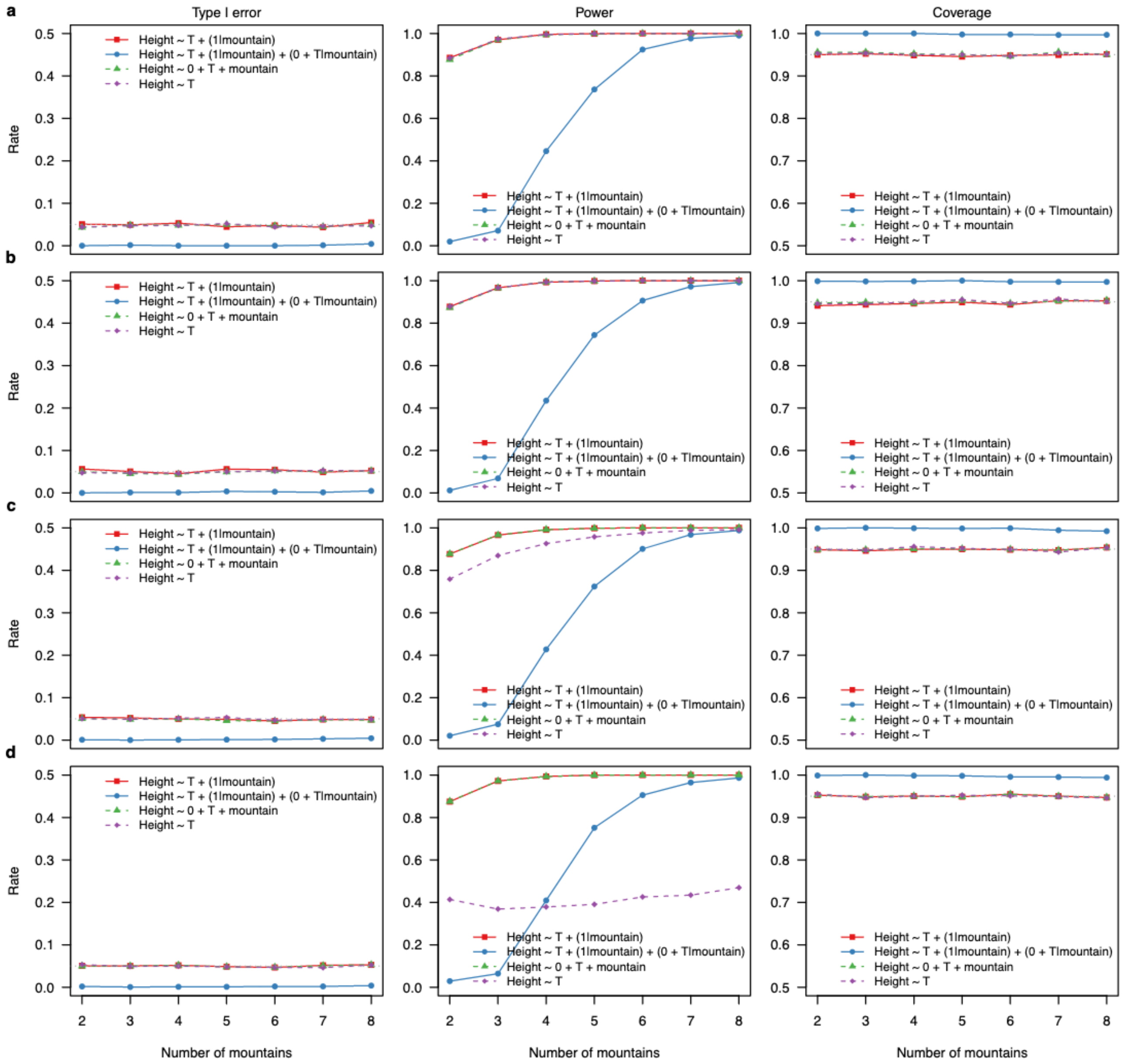


**Figure S12:** Type I error rates, power, and coverage for linear (mixed effect) models fitted with *lme4* package to simulated data with 2-8 mountains for scenario A (random intercept for each mountain) and 25 observations per mountain range. a-d show the results for different standard deviations of the random effect (0.01,0.1, 0.5 and 2.0). For each scenario, 5,000 simulations and models were tested.

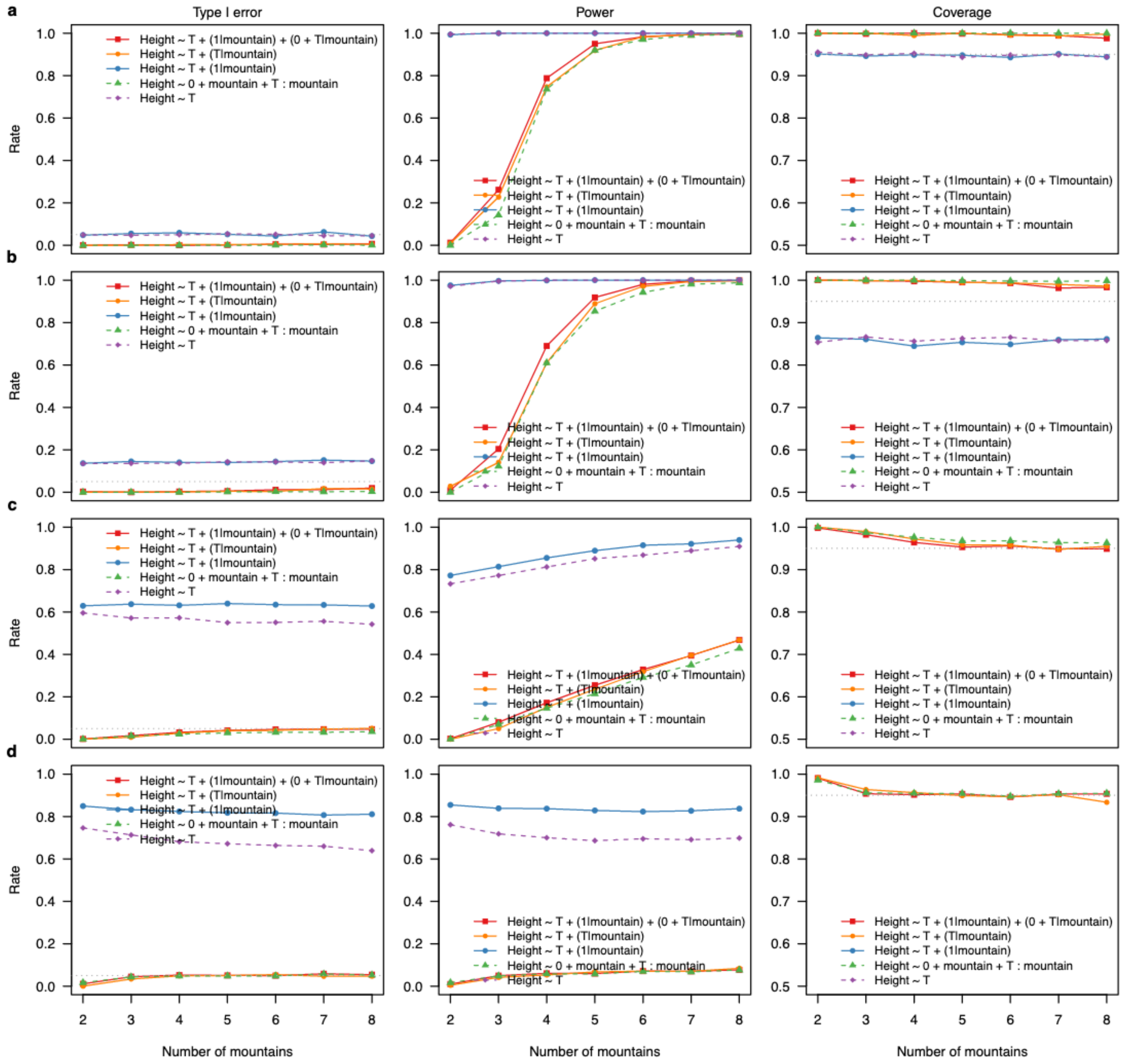


**Figure S13:** Type I error rates, power, and coverage for linear (mixed effect) models fitted with lme4 to simulated data with 2-8 mountains for scenario B (random intercept and random slope for each mountain) and on average 50 observations per mountain range. a-d show the results for different standard deviations (0.01,0.1, 0.5 and 2.0). For each scenario, 5,000 simulations and models were tested.

### 4. Influence of sample size on statistical properties

Generalized linear models (mixed or not) with non-gaussian distributions require much more data to have a high probability to truly detect a significant effect (power). We ran additional data with different sample sizes per level in the grouping variable (25, 50, 100, 200) and compared type I, power, and coverage for the generalized linear models (hypothesis 2, Box1) in both scenarios A (Fig. S14) and B (Fig. S16), as well as the for the linear (mixed-effect) models for scenario B (Fig. S15).

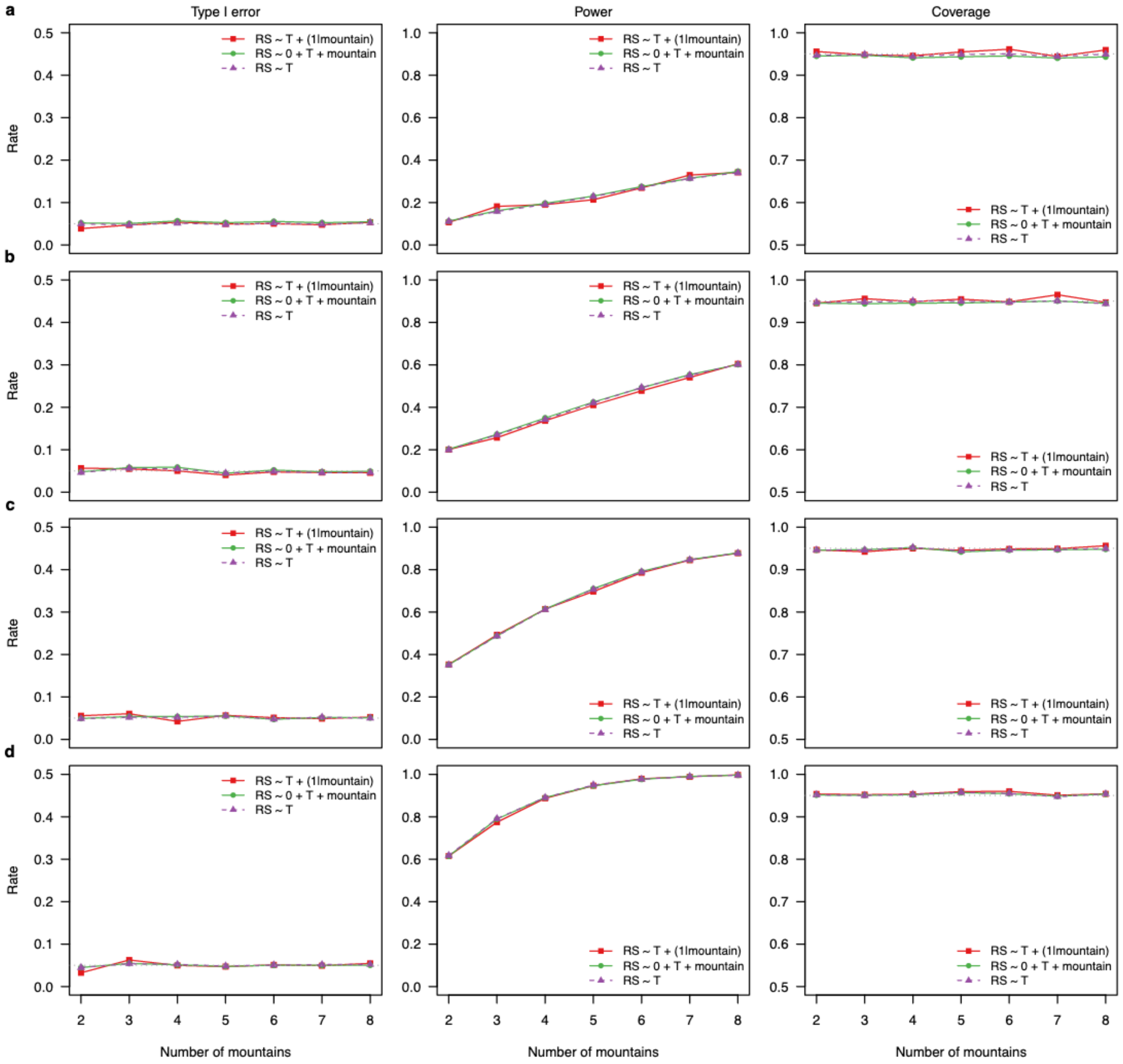


**Figure S14:** Type I error rates, power, and coverage for generalized linear (mixed effect) models fitted with *lme4* package to simulated data with 2-8 mountains for scenario A (random intercept for each mountain). a-d show the results for different numbers of observations for each mountain (25, 50, 100, and 200). For each scenario, 5,000 simulations and models were tested.

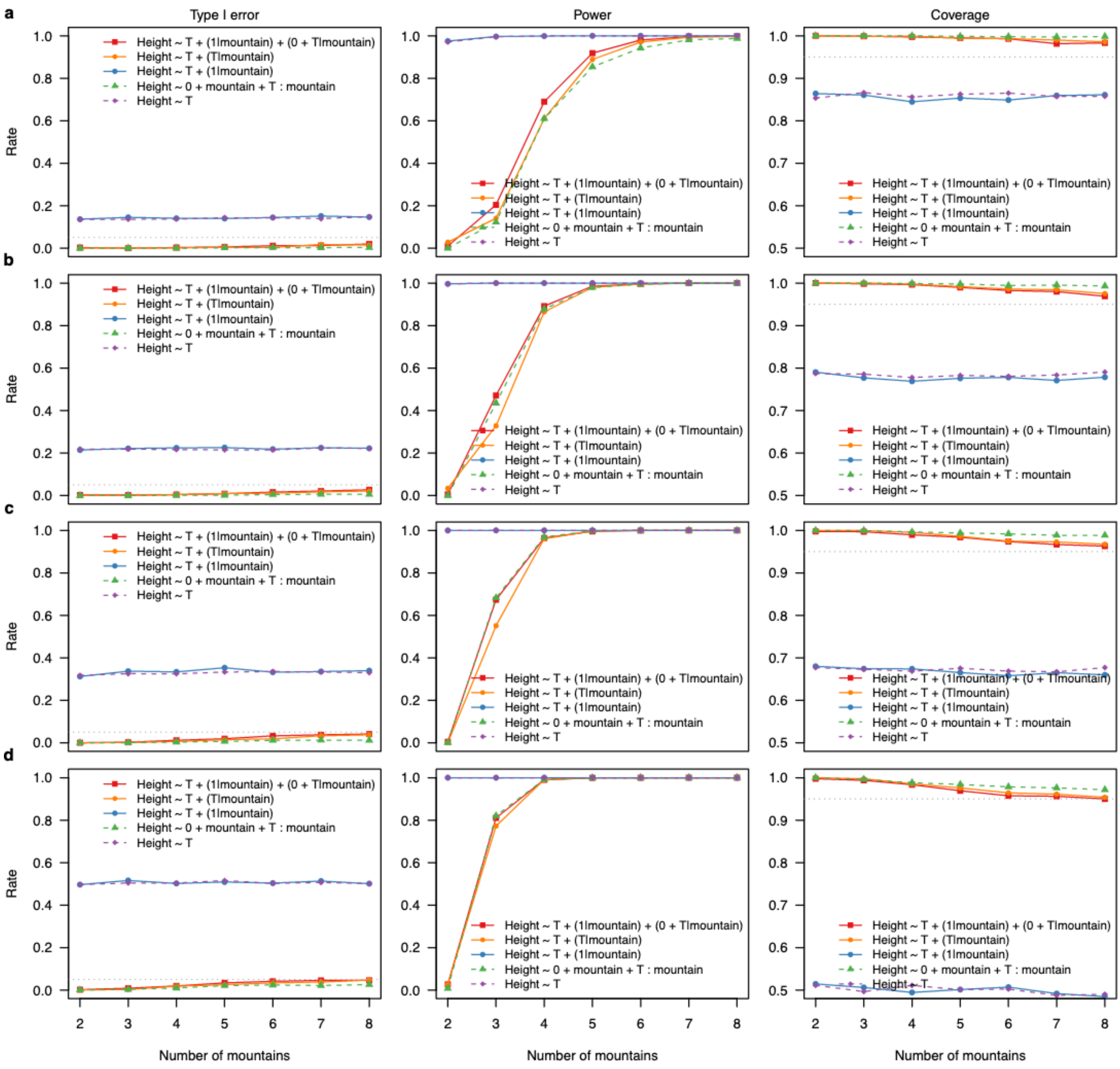


**Figure S15:** Type I error rates, power, and coverage for linear (mixed effect) models fitted with *lme4* package to simulated data with 2-8 mountains for scenario B (random intercept and slope for each mountain). a-d show the results for different numbers of observations for each mountain (50, 100, 200, and 500). For each scenario, 5,000 simulations and models were tested.

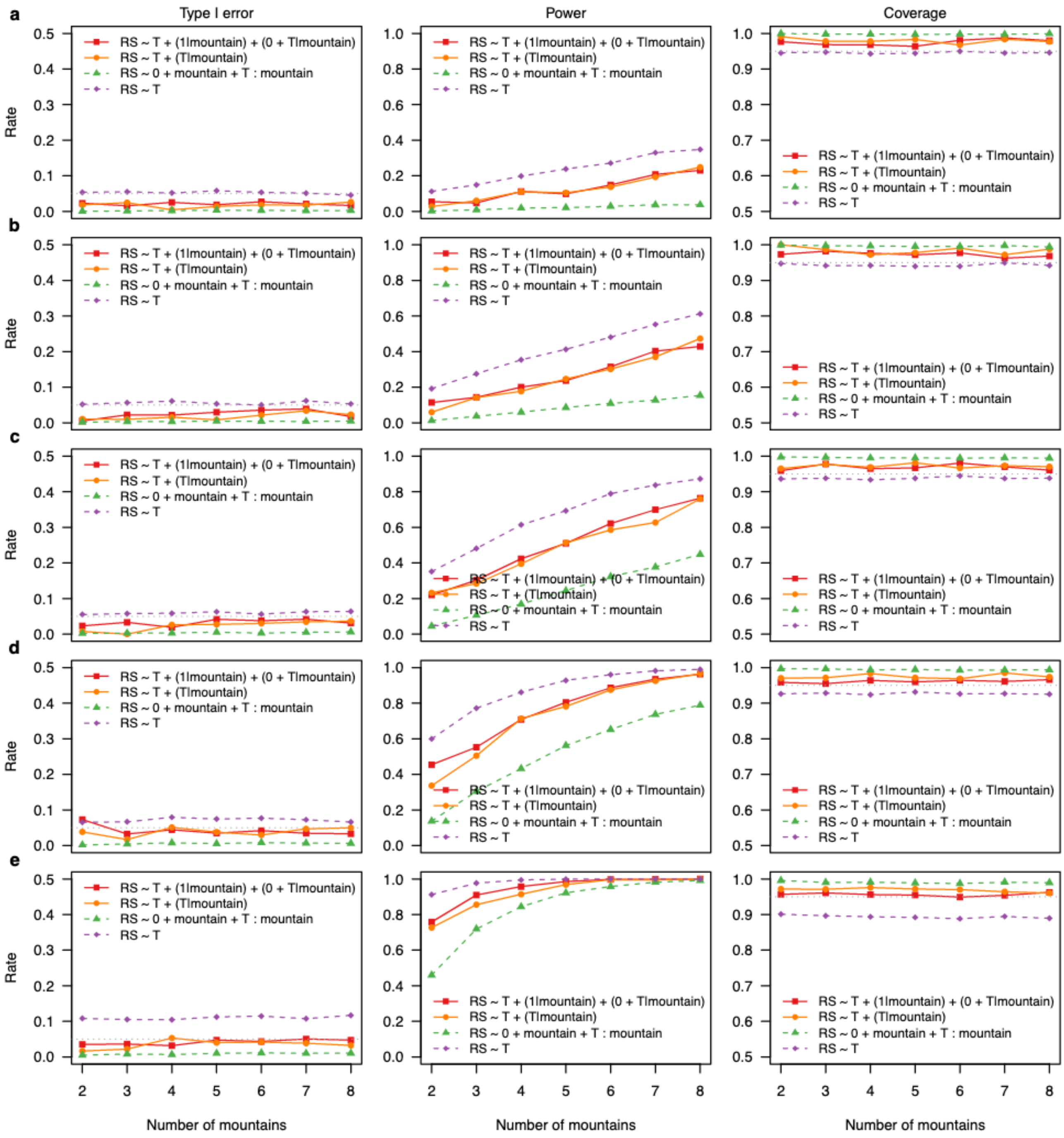


**Figure S16:** Type I error rates, power, and coverage for generalized linear (mixed effect) models fitted with *lme4* package to simulated data with 2-8 mountains for scenario B (random intercept and slope for each). a-e show the results for different numbers of observations for each mountain (25, 50, 100, 200, and 500). For each scenario, 5,000 simulations and models were tested.

### 5. Calculation of the mean temperature effect in fixed-effect models with interaction

As fixed-effect models with interactions estimate the effect of one level and its contrasts to the other levels, the population effect of temperature itself is not estimated in the R default parametrization. To calculate the population effect and its significance to be able to compare it to mixed-effect models results, we estimate the grand mean and its standard error via bootstrapping. To do so, we sample from the multivariate normal distribution reported from the fitted linear for the interactions. Let S be a big enough sample from this distribution, then the mean of S is the grand mean temperature effect, and the standard error of S is the standard error of the mean. In R language we can calculate p-values using 2000 samples given the effect estimates *effects* and the covariance matrix of the individual level effects *V* from the linear model in the following way:

```
S = mvtnorm::rmvnorm(2000, mean = effects, sigma = V)
eff = mean(S)
se = sd(S)/sqrt(mountain-1)
```

We then used t-values for LMs and z-values for GLMs for a better comparison with the respective packages to calculate p-values.

To check whether our calculation of the grand mean behaves approximatively correctly, i.e. with no effect we would expect a uniform distribution of the p-values, we simulated scenario B with no grand mean (effect was set to 0), different standard deviations of the varying random intercept and slope (0.1, 0.5, and 1.0), and unbalanced (30 observations for each group but 1000 observations for the last one) and balanced (30 observations for each group) study design. We found that the distribution of p-values is approximatively correct for larger standard deviations (which is linked to the number of observations in each group, Fig. S17).

We repeated the simulations for GLMs and we found that the distributions of p-values were on average left-skewed (Fig. S18). The nature of the logit link in the GLM explains the increase in left skewness with higher standard deviations of the random effects (Fig. S18).

Using zero sum contrasts is another way to calculate the average affects over a grouping variable. We repeated the above-mentioned simulation to compare the zero-sum contrasts method to our bootstrapping approach. However, we found highly right-skewed distribution of p-values for the zero-sum contrasts for both models, the LM and the GLM (Fig. S19, S20). Thus, the zero-sum contrasts method leads to high type I errors and very high power for LMs and GLMs, while our bootstrapping method leads to correct type I errors and power for LMs, but higher than expected type I error rates and less power for the GLMs. Therefore, we think that our bootstrapping approach is preferable over the zero-sum contrast method.

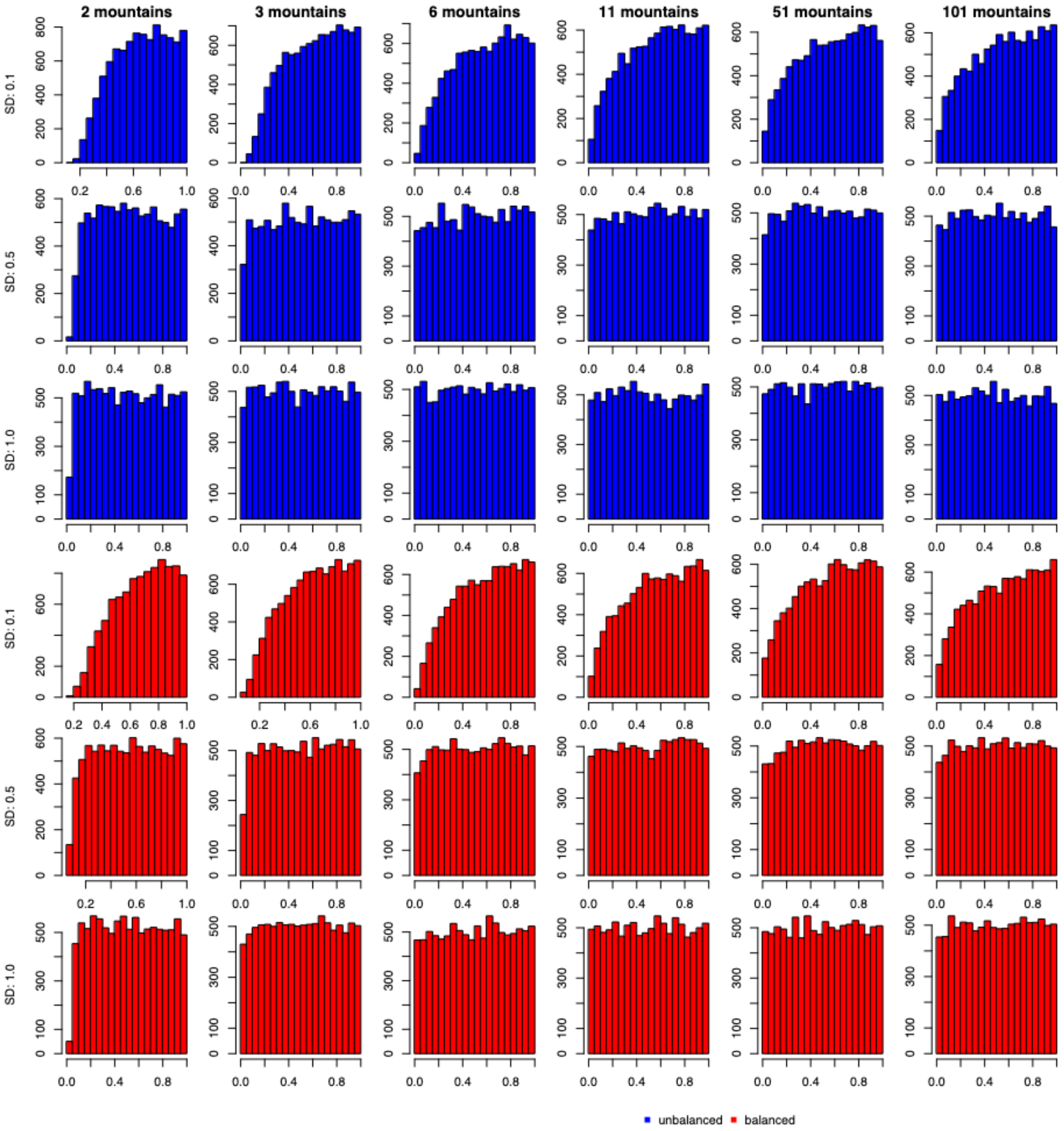


**Figure S17:** The distribution of p-values (based on t-statistics) for the grand mean calculation of linear models for different simulation scenarios: 2 – 101 mountains, different SDs for the random effects (0.1, 0.5, and 1.0), and for unbalanced (blue) and balanced (red) study design. Each scenario was simulated 10,000 times.

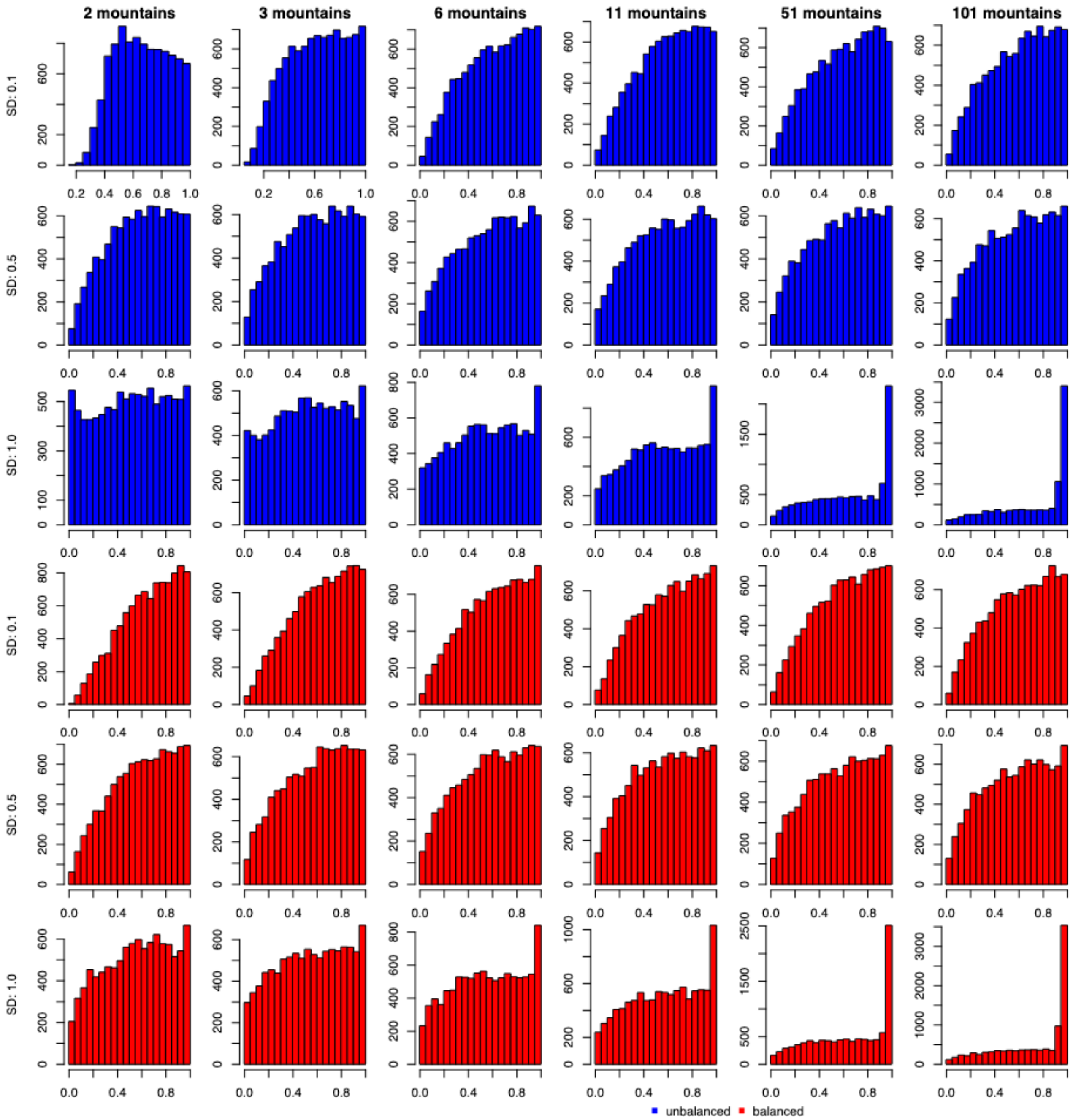


**Figure S18:** The distribution of p-values (based on z-statistics) for the grand mean calculation of generalized linear models for different simulation scenarios: 2 – 101 mountains, different SDs for the random effects (0.1, 0.5, and 1.0), and for unbalanced (blue) and balanced (red) study design. Each scenario was simulated 10,000 times.

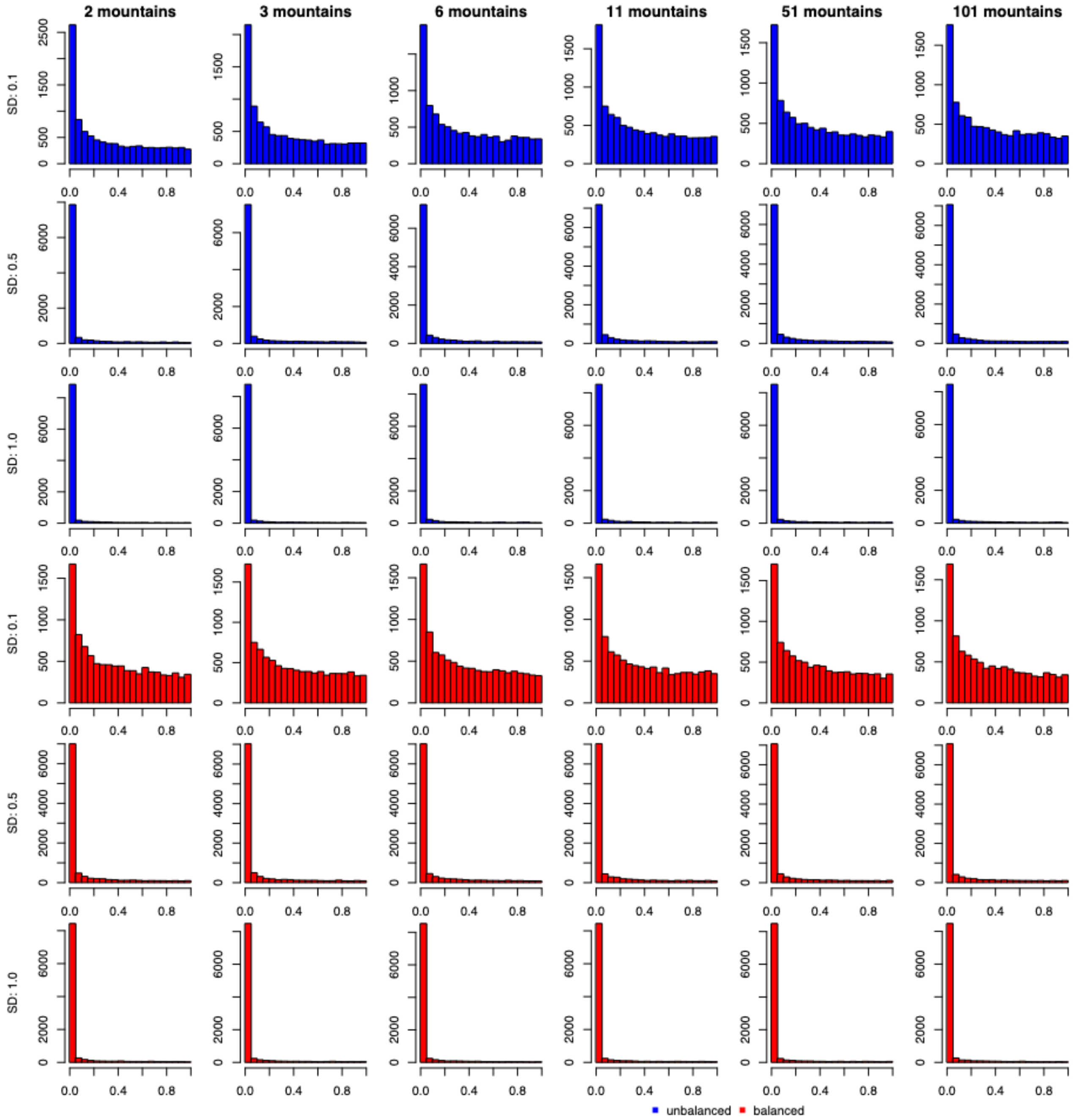


**Figure S19:** The distribution of p-values (based on t-statistics) for the grand mean calculation in linear models (based on zero sum contrasts) for different simulation scenarios: 2 – 101 mountains, different SDs for the random effects (0.1, 0.5, and 1.0), and for unbalanced (blue) and balanced (red) study design. Each scenario was simulated 10,000 times.

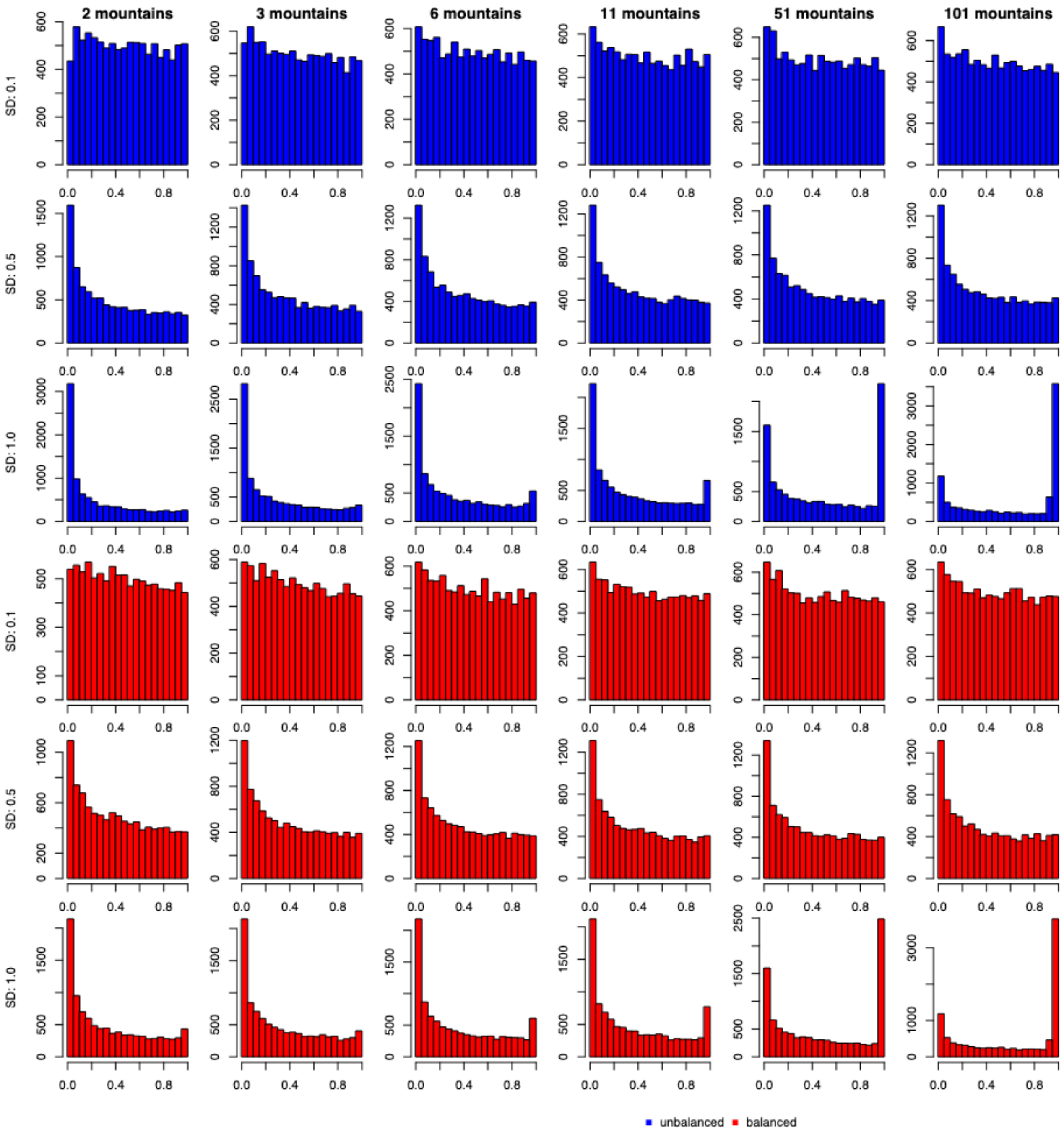


**Figure S20:** The distribution of p-values (based on t-statistics) for the grand mean calculation in generalized models (based on zero sum contrasts) for different simulation scenarios: 2 – 101 mountains, different SDs for the random effects (0.1, 0.5, and 1.0), and for unbalanced (blue) and balanced (red) study design. Each scenario was simulated 10,000 times.

### 6. Distribution of population effect estimates

To check what causes different type I error rates, we plotted the distribution of estimated population mean effects for scenario B for a different number of mountains (2, 3, 5, and 7, corresponding to panels a-d in Fig. S21 and S22) for the intercept (Fig. S21) as well as for the slope (Fig. S22). We found no differences for the different models for the estimates of the population effect (Fig. S21, 22). We found that with increasing number of mountains the variance of the distribution declines.

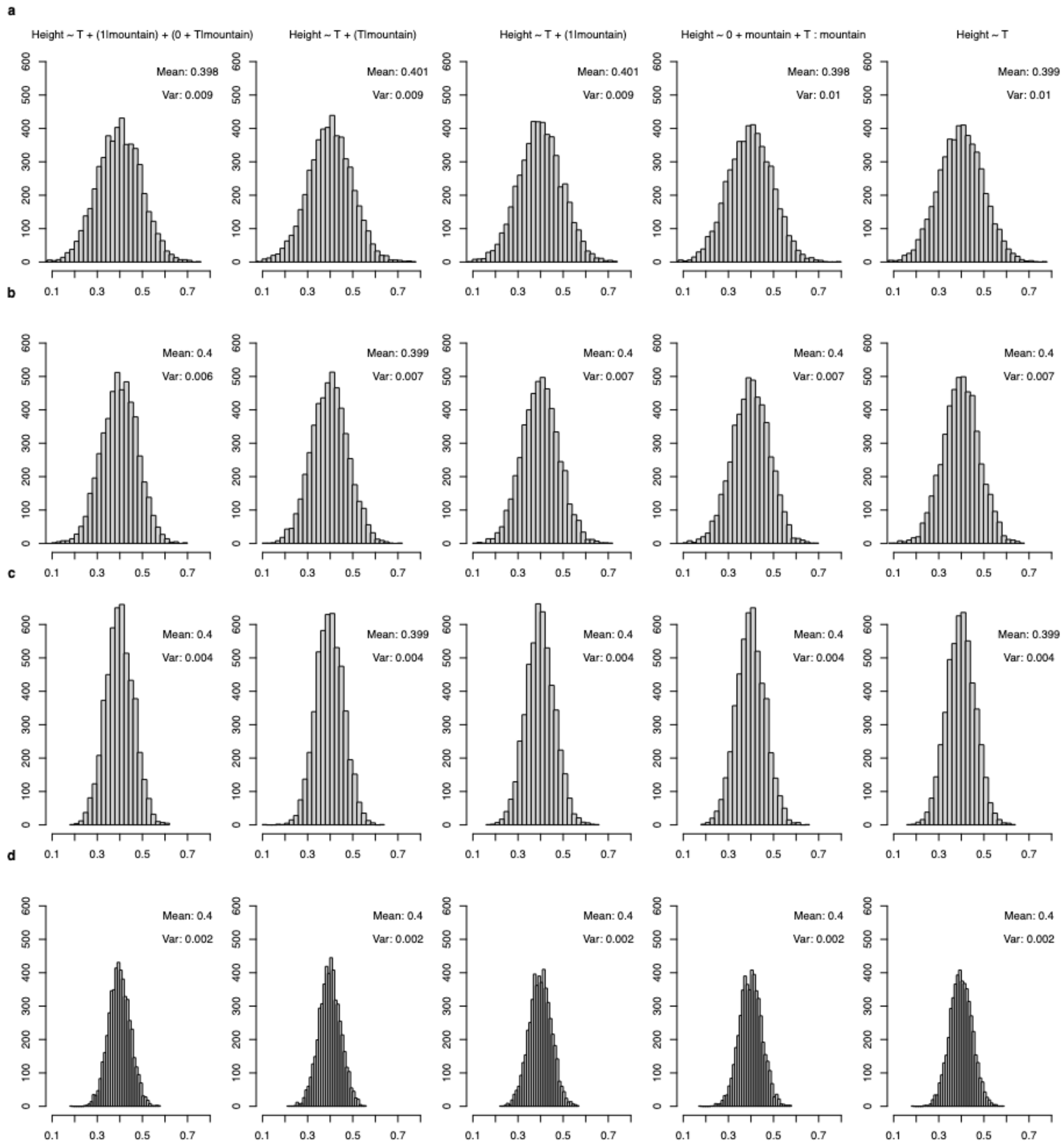


**Figure S21:** The distribution of the population slope effect (grand mean, ecological effect) of temperature for scenario B. a-d show different number of mountains (2, 3, 5, and 7) and each column corresponds for a different model which we tested for scenario B (see Figure 2 in main text).

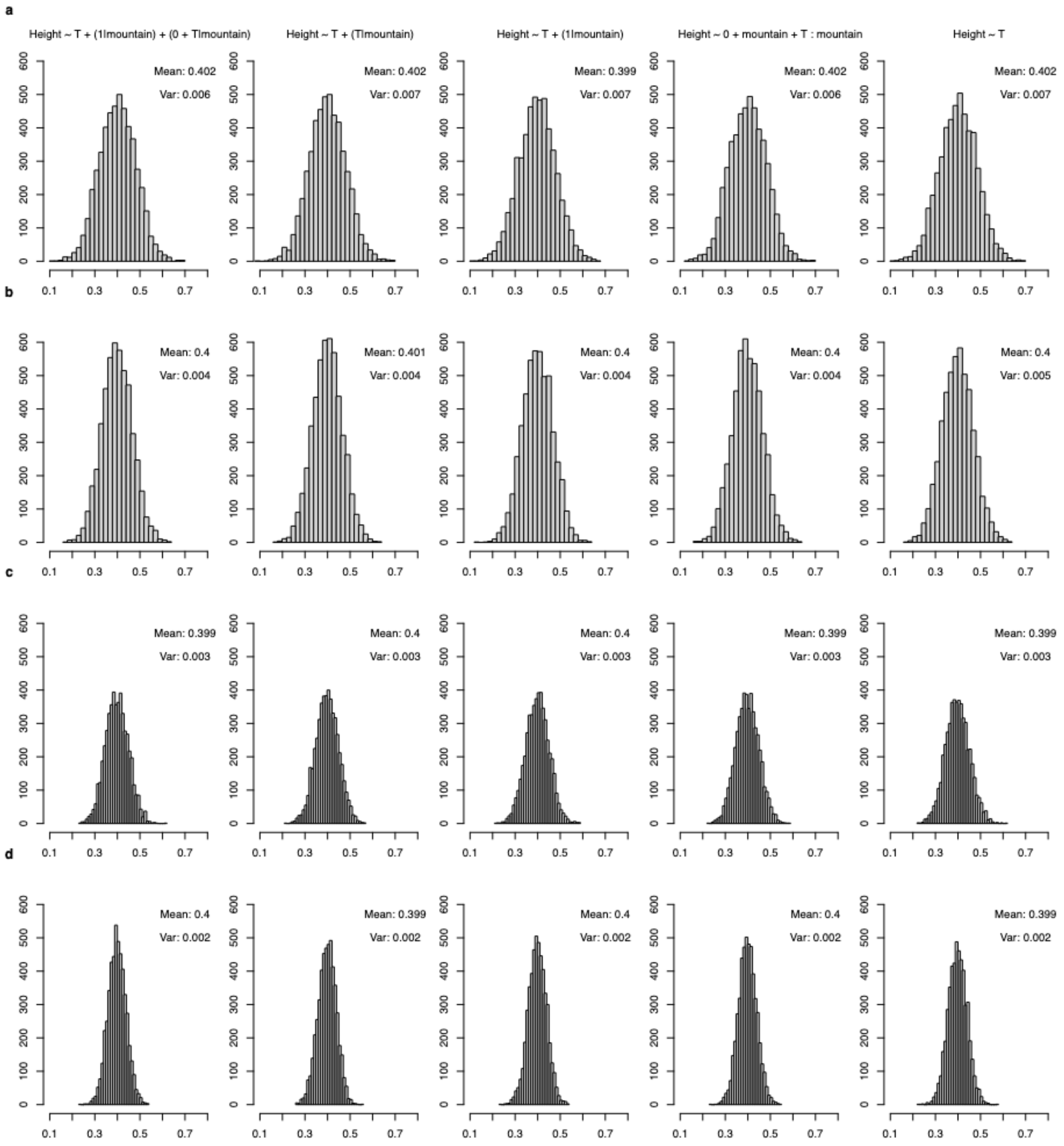


**Figure S22:** The distribution of the population intercept effect for scenario B. a-d show different number of mountains (2, 3, 5, and 7) and each column corresponds for a different model which we tested for scenario B (see Figure 2 in main text).
